# Gene regulatory Networks Reveal Sex Difference in Lung Adenocarcinoma

**DOI:** 10.1101/2023.09.22.559001

**Authors:** Enakshi Saha, Marouen Ben Guebila, Viola Fanfani, Jonas Fischer, Katherine H. Shutta, Panagiotis Mandros, Dawn L. DeMeo, John Quackenbush, Camila M. Lopes-Ramos

**Author notes:** **Corresponding Author:** Camila M Lopes-Ramos.

## Abstract

Lung adenocarcinoma (LUAD) has been observed to have significant sex differences in incidence, prognosis, and response to therapy. However, the molecular mechanisms responsible for these disparities have not been investigated extensively. Sample-specific gene regulatory network methods were used to analyze RNA sequencing data from non-cancerous human lung samples from The Genotype Tissue Expression Project (GTEx) and lung adenocarcinoma primary tumor samples from The Cancer Genome Atlas (TCGA); results were validated on independent data. We observe that genes associated with key biological pathways including cell proliferation, immune response and drug metabolism are differentially regulated between males and females in both healthy lung tissue, as well as in tumor, and that these regulatory differences are further perturbed by tobacco smoking. We also uncovered significant sex bias in transcription factor targeting patterns of clinically actionable oncogenes and tumor suppressor genes, including *AKT2* and *KRAS*. Using differentially regulated genes between healthy and tumor samples in conjunction with a drug repurposing tool, we identified several small-molecule drugs that might have sex-biased efficacy as cancer therapeutics and further validated this observation using an independent cell line database. These findings underscore the importance of including sex as a biological variable and considering gene regulatory processes in developing strategies for disease prevention and management.

## Introduction

Lung adenocarcinoma (LUAD) exhibits significant sex differences in incidence, prognosis and response to therapy. LUAD has been observed to be more prevalent in females than males [1, 2, 3], with the sex difference being more pronounced among nonsmokers [4]. However, males with LUAD have more severe disease and poorer survival outcomes compared to females with the disease [5]. Treatment responses and toxicity are also influenced by sex [5]; while females usually respond better to chemotherapy compared to males [6], immune checkpoint inhibitors have been found to be more effective in males [7] with lung cancer.

Increased susceptibility of LUAD in females may partially be attributed to the effect of estrogens on lung carcinogen metabolism. For example, polymorphisms in cytochrome P450 1A1 (*CYP1A1*) and glutathione *S*-transferase M1 (*GSTM1*) may contribute to the increased risk of females for lung cancer. Females with the *CYP1A1* mutant/*GSTM1* null genotypes face an elevated risk, regardless of their smoking history, potentially influenced by estrogen exposure [8]. Hormonal influences could contribute not only to lung cancer incidence, but also its development and survival outcomes [9]. Prior research has detected the existence of estrogen receptors in malignant lung tissues in both sexes [10]. However, the effects of sex steroid hormones may not account for all differences in how males and females respond to environmental carcinogens including smoking [4]. Among other factors, higher DNA adduct levels and more frequent mutations in the proto-oncogene *KRAS* in females have also been cited as a possible contributor governing higher lung cancer risk in females [11]. Genetic and metabolic factors have also been cited as potential mediators for the better prognostic outcomes in females [12, 13] compared to males with lung cancer. While previous studies have focused on molecular alterations and gene expression alone [14, 15], an integrative analysis of multi-omics data from a systems perspective can offer valuable insights into sex-specific regulatory mechanisms linked to both lung cancer incidence and clinical outcome.

Despite documented sex differences in LUAD risk and subsequent disease outcome, most methods used in the development and selection of cancer therapeutics do not consider biological sex differences, in part because their molecular drivers are poorly understood, and partly because clinical trials are not designed to address sex-specific effects. Understanding the regulatory processes that differentiate between the sexes in both healthy lung tissue and in LUAD will not only help to elucidate disease mechanisms but also identify more effective therapeutic approaches for both sexes.

We inferred gene regulatory networks using PANDA [16] and LIONESS [17], methods that in combination integrate genome-wide transcription factor binding site maps, transcription factor protein-protein interaction data, and gene expression profiles to produce sample-specific regulatory network models that have successfully uncovered sex-specific regulatory drivers of health and disease in previous studies [18, 19, 20, 21]. We compared these sample-specific regulatory networks between males and females to identify genes and biological pathways targeted by transcription factors in a sex-biased manner in both healthy lung tissue and in LUAD samples. We further explored how this sex bias is influenced by smoking behavior, a significant risk factor for lung cancer.

As a primary measure of regulatory network differences, we used differential gene targeting, which identifies significant changes in the network model transcription factor repertoire controlling each gene. Among healthy samples, genes associated with cell adhesion and cell proliferation were highly targeted among female nonsmokers, while in tumor samples these genes showed higher targeting in males, irrespective of smoking history. Genes associated with immune pathways exhibited higher targeting in tumor samples from females than in those from males, suggesting the potential for sex-based differential response to cancer immunotherapy. Pathways with known relevance in chemotherapy response such as drug metabolism cytochrome P450 (CYP450) showed higher targeting in females, compared to males. Furthermore, an elevated targeting of drug metabolism CYP450 was also associated to favorable survival outcomes in response to chemotherapy among females but not males. We also uncovered significant sex bias in transcription factor targeting of oncogenes and tumor suppressor genes, including *AKT2* and *KRAS* that suggests lung cancer drugs targeting these genes might exhibit differences between the sexes in both efficacy and toxicity. Using an *in-silico* drug repurposing tool, we identified several small-molecule drugs that might have sex-biased efficacy as cancer therapeutics and further validated this hypothesis using an independent cell line database.

## Results

### The Role of Differential Gene Regulation by Sex in Incidence Risk of LUAD

To understand why females have a higher risk of developing LUAD compared to males, especially among nonsmokers, we compared male and female gene regulatory networks inferred from GTEx for healthy lung samples [**Figure 2**]. We identified several key pathways that are targeted by transcription factors in a sex-biased manner in healthy lung that shed light on potential mechanisms driving sex difference in disease risk. For nonsmokers we observed increased targeting in females compared to males (FDR<0.05) of pathways responsible for cell proliferation, cell adhesion and migration, including the hedgehog signaling pathway, WNT signaling pathway, notch signaling pathway, ERBB signaling pathway, non-small cell lung cancer, focal adhesion and adherens junction [**Figure 2**]. We validated these findings in healthy lung samples from an independent dataset (LGRC) [**Figure D.2**]. For smokers, all these pathways mentioned were more highly targeted in males than females, however, in the LGRC dataset we were only able to validate this finding for pathways associated with non-small cell lung cancer and hedgehog signaling [**Figure D.2**].

**Figure 1:**
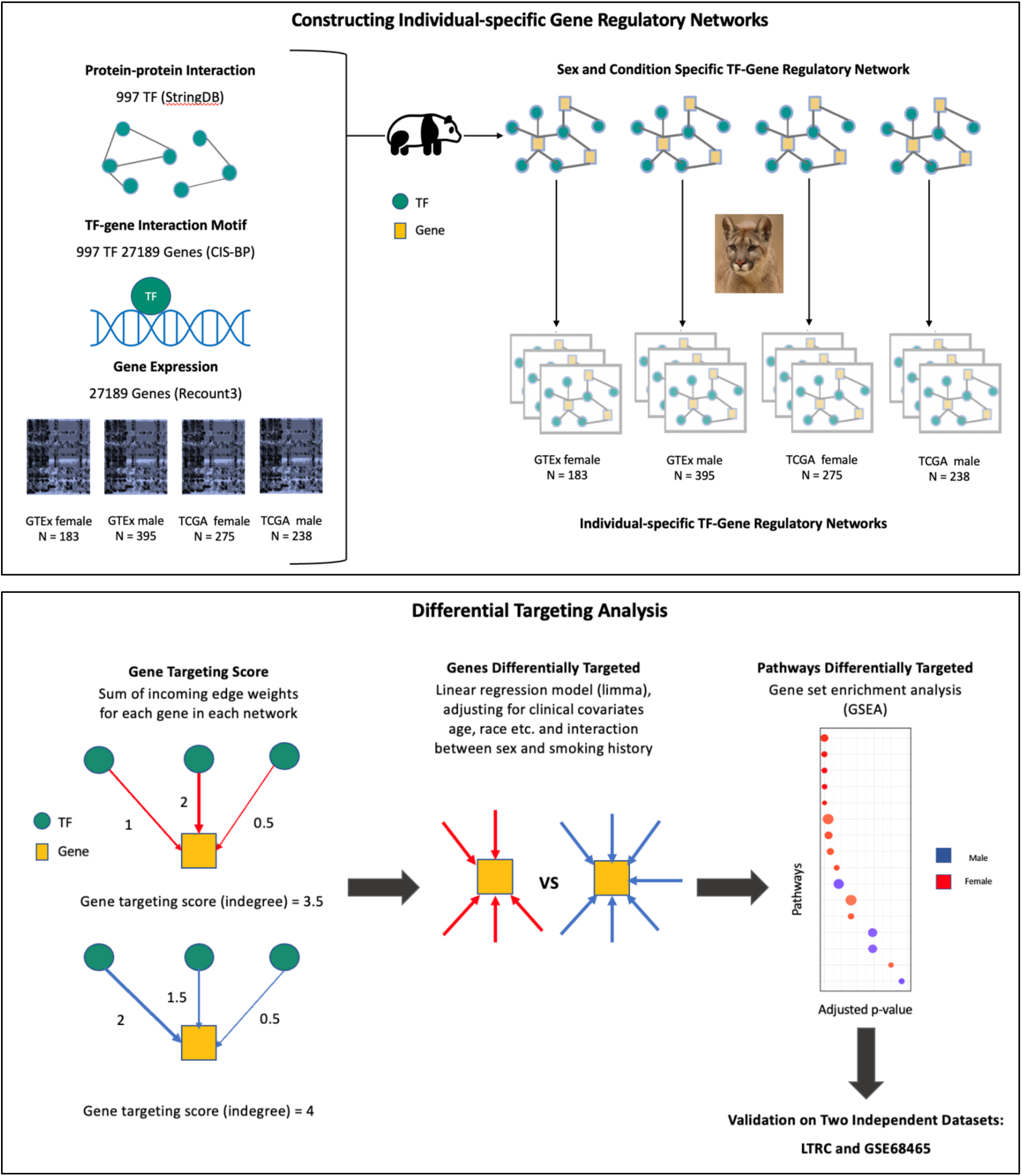
Schematic overview of the study. Top box, overview of the approach used to construct individual specific gene regulatory networks with PANDA and LIONESS by integrating information on protein-protein interaction between transcription factors (TFs), TF-gene motif binding, and gene expression data of GTEx healthy lung tissues and TCGA lung adenocarcinoma (LUAD) primary tumor samples from Recount3. Bottom box, overview of the differential targeting analysis and independent datasets for validation.

**Figure 2:**
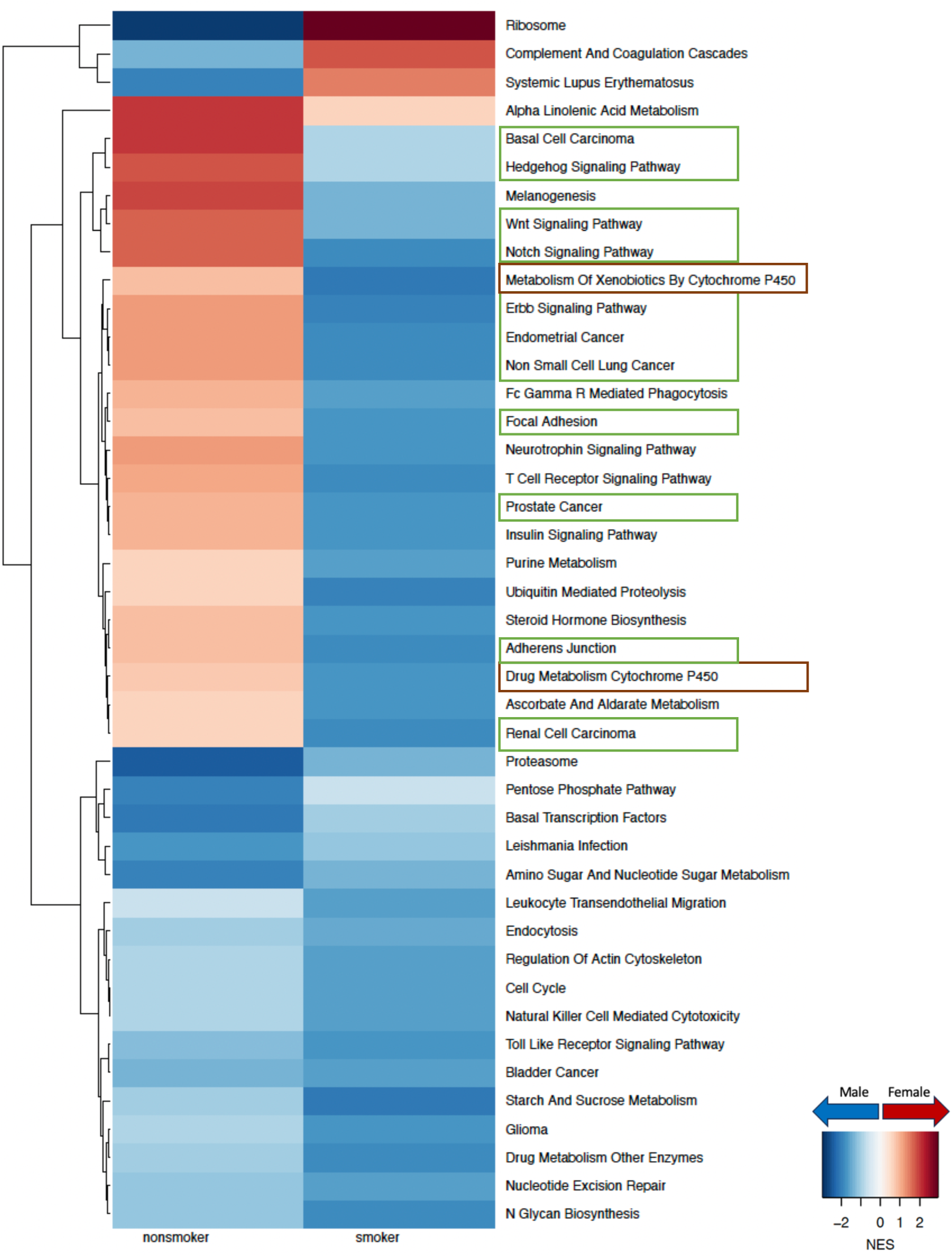
Sex difference in GTEx healthy lung samples within nonsmokers and smokers. Normalized enrichment scores (NES) from gene set enrichment analysis (GSEA) using KEGG pathways are shown for all pathways that have significant (adjusted p-value < 0.05) sex difference among either nonsmokers or smokers. Pathways with higher targeting in male are marked blue and pathways with higher targeting in female are marked red. Green boxes highlight pathways associated with cell proliferation and brown boxes highlight pathways associated with environmental carcinogen metabolism.

The CYP450 drug metabolism pathway, which is associated with environmental carcinogen metabolism [11] also had higher targeting in female among nonsmokers and in male among smokers, within both GTEx [**Figure 2**] and LGRC [**Figure D.2**] control samples.

Based on our analysis we observe that in healthy human lung, pathways related to cell proliferation and environmental carcinogen metabolism were differentially regulated between males and females which might contribute to the difference in risk of developing LUAD between the sexes.

### Understanding Sex Difference in LUAD Prognosis through Differential Gene Regulation

To understand why males have poorer prognosis than females with LUAD, we compared the gene regulatory networks of primary tumors from males and females from the TCGA and identified key pathways with a sex-biased targeting pattern by transcription factors. Specifically, we found that pathways involved in cell adhesion, cell proliferation, and cell migration, such as WNT signaling pathway, pathways in cancer, tight junction, and adherens junction, all have higher targeting in tumors from males compared to those from females irrespective of smoking status. It is interesting to note that for nonsmokers [**Figure 3**], cell proliferation and migration-related pathways switched from having higher targeting in healthy females to having higher targeting in male tumors. And for smokers [**Figure 4**], pathways related to cell proliferation and cell migration that were already highly targeted in healthy males become even more highly targeted in male tumors, compared to females.

**Figure 3:**
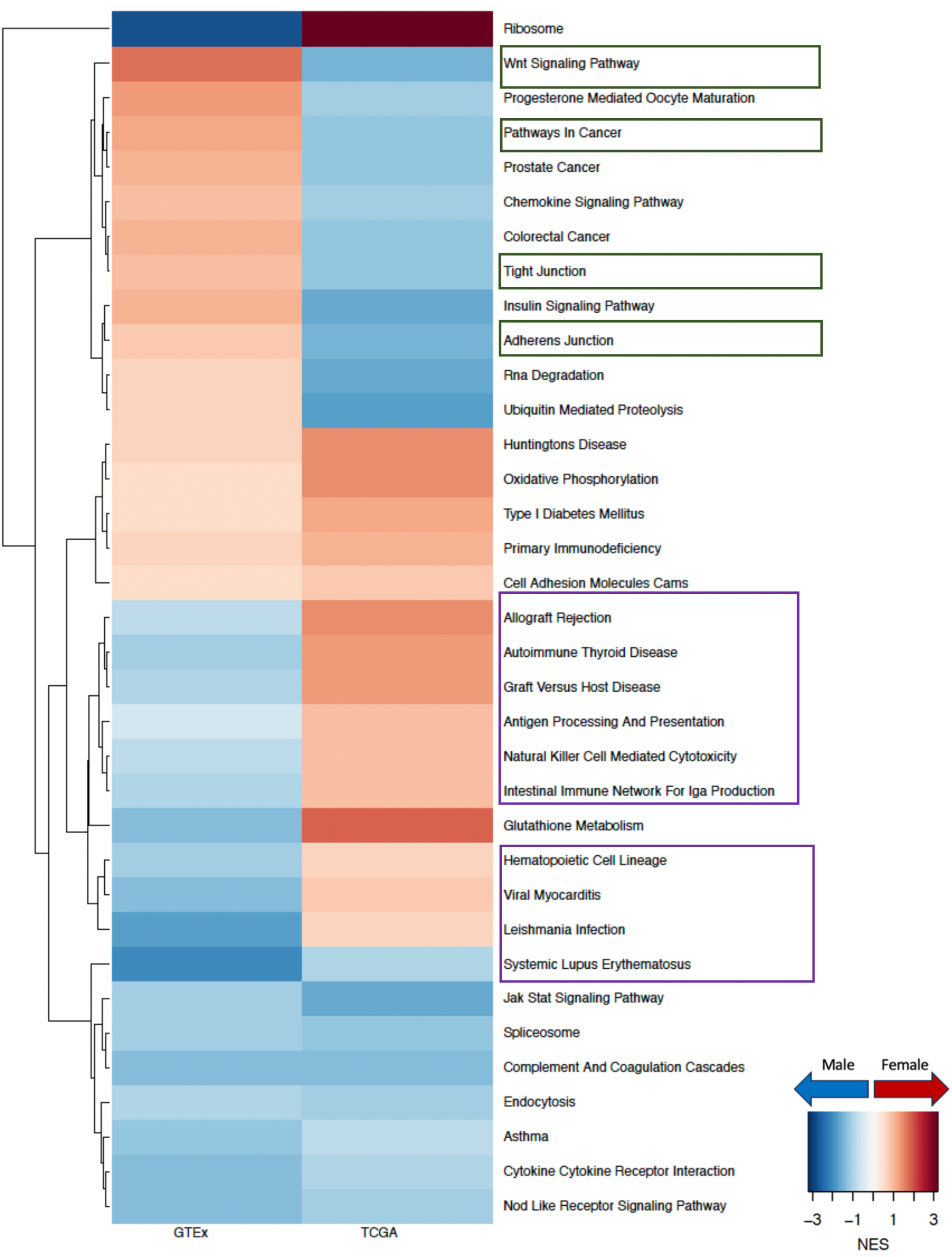
Sex difference among nonsmokers in GTEx healthy lung and in TCGA LUAD. Normalized enrichment scores (NES) from GSEA using KEGG pathways are shown for all pathways that have significant (adjusted p-value < 0.05) sex difference among *either TCGA nonsmokers or TCGA smokers*. Pathways with higher targeting in male are marked blue and pathways with higher targeting in female are marked red. Green boxes highlight pathways associated with cell proliferation and purple boxes highlight pathways associated with immune response.

**Figure 4:**
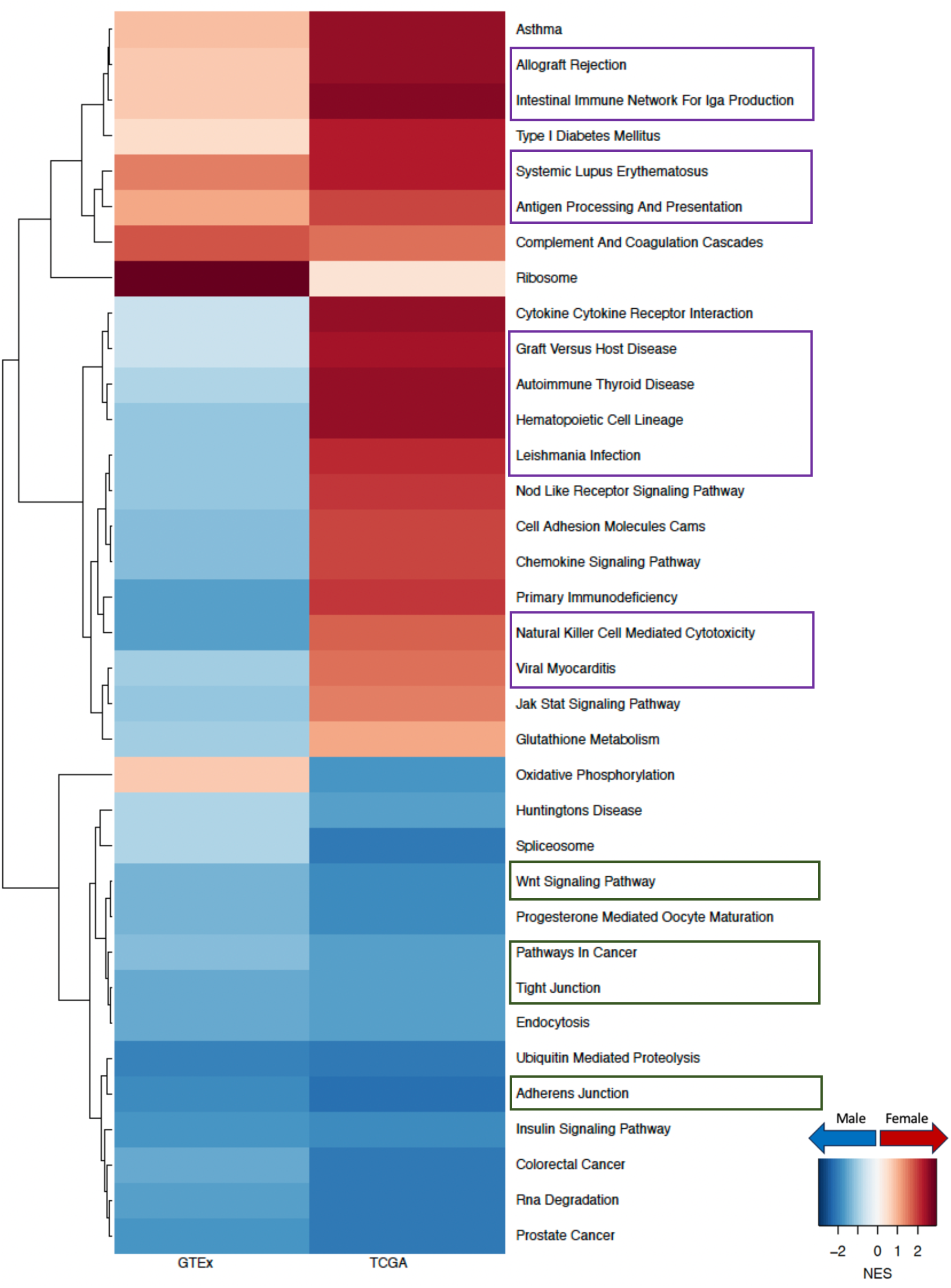
Sex difference among smokers in GTEx healthy lung and in TCGA LUAD. Normalized enrichment scores (NES) from GSEA using KEGG pathways are shown for all pathways that have significant (adjusted p-value < 0.05) sex difference among *either TCGA nonsmokers or TCGA smokers*. Pathways with higher targeting in male are marked blue and pathways with higher targeting in female are marked red. Green boxes highlight pathways associated with cell proliferation and purple boxes highlight pathways associated with immune response.

We replicated our network analysis using an independent LUAD dataset (GSE68465) [**Figure D.3**] and validated that among nonsmokers, WNT signaling pathway and tight junction were more highly targeted in male tumors than in those from females. We also validated that among smokers, pathways in cancer and adherens junction showed higher targeting among male tumors, consistent with the results from TCGA.

We then turned our attention to oncogenes and tumor suppressor genes cataloged in the COSMIC database [22] and found these to also be highly differentially targeted between the sexes in both healthy and tumor samples [**Figure 5**]. Among nonsmokers in healthy GTEx lung samples, both oncogenes and tumor suppressor genes showed higher targeting (p-value of Wilcoxon signed rank test is 2.229e-09 for oncogenes and 3.614e-05 for tumor suppressor genes) in females compared to males. Whereas among the nonsmokers in the TCGA tumor samples, both oncogenes and tumor suppressor genes showed higher targeting in male samples (p-value of Wilcoxon signed rank test is 2.334e-09 for oncogenes and 5.217e-07 for tumor suppressor genes), which may help explain poorer prognosis in males compared to females. For smokers, oncogenes and tumor suppressor genes showed higher targeting for males than females in both healthy lung samples from GTEx (p-value of Wilcoxon signed rank test is 3.546e-08 for oncogenes and 2.296e-12 for tumor suppressor genes), as well as LUAD tumors from TCGA (p-value is 5.906e-08 for oncogenes and 2.296e-12 for tumor suppressor genes).

**Figure 5:**
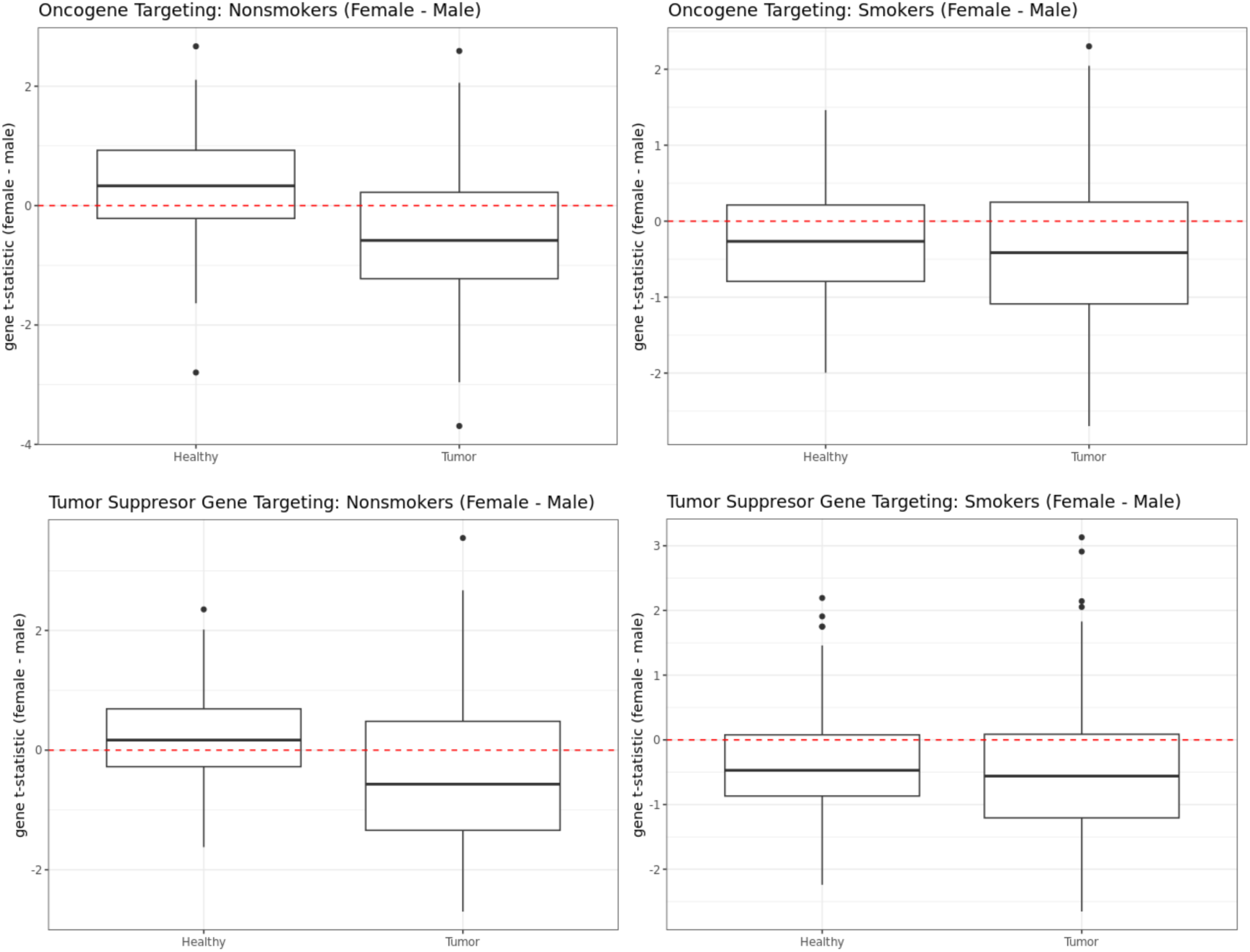
Sex difference in targeting of oncogenes (top row) and tumor suppressor genes (bottom row) in GTEx and TCGA nonsmokers (left column) and smokers (right column).

To understand whether sex differences in regulation of biological pathways might explain poorer survival among males with LUAD, we performed survival analysis on TCGA data using a Cox proportional hazard model for each of these pathways. We found a higher targeting of the RNA Degradation pathway to be associated with poorer survival outcome in males (z-score of the coefficient corresponding to pathway score is 2.030 with p-value 0.042) but did not have any impact in females (z-score of the coefficient corresponding to pathway score is -0.740 with p-value 0.459). The leading genes contributing towards a higher targeting of RNA degradation among males include *CNOT1* [23], *CNOT2* [24], *CNOT3* [25] and *DCP1A* [26], all of which have previously been found to have prognostic significance in various cancers, including non-small cell lung cancer.

### Sex Difference in Immunotherapy

Among GTEx smokers [**Figure 4**], immune-related pathways such as allograft rejection, intestinal immune response for IGA production, systemic lupus erythematosus, antigen processing and presentation, all showed higher targeting in females. This female bias was even more strongly evident in TCGA tumor samples from smokers. Among GTEx nonsmokers [**Figure 3**] these pathways were more highly targeted in males. However, in TCGA we found higher targeting in females—except for systemic lupus erythematosus which remained highly targeted in male tumor samples—but the observed sex difference was smaller than that in GTEx. Other immune pathways such as hematopoietic cell lineage and natural killer cell mediated cytotoxicity showed higher targeting in male within GTEx and switched to higher targeting in female within TCGA, irrespective of smoking status. This pattern of female-bias in targeting of immune pathways was validated in tumor samples from GSE68465 among smokers.

We performed immune cell type deconvolution analysis of TCGA data and found that, consistent with a higher targeting of immune pathways in females, various immune cell proportions including natural killer cells, CD4+ naive T cells, myeloid dendritic cells and B cells were higher among female tumor samples than male tumor samples [**Figure 6**]. The only exceptions are CD4+ Th2 helper cells that are present in higher proportions among male samples. Differential targeting of immune pathways, along with a sex-biased infiltration of immune cells, might contribute to varying degrees of efficacy of immune checkpoint inhibitors shown to exist among males and females with LUAD [**Table C.1**] [27]. Interestingly, within healthy samples from GTEx, we did not find any sex difference in the proportion of immune cells that had sex-biased infiltration rate in TCGA [**Figure D.4**], the only exception being natural killer T cells, which showed higher proportion in males compared to females among nonsmokers.

**Figure 6:**
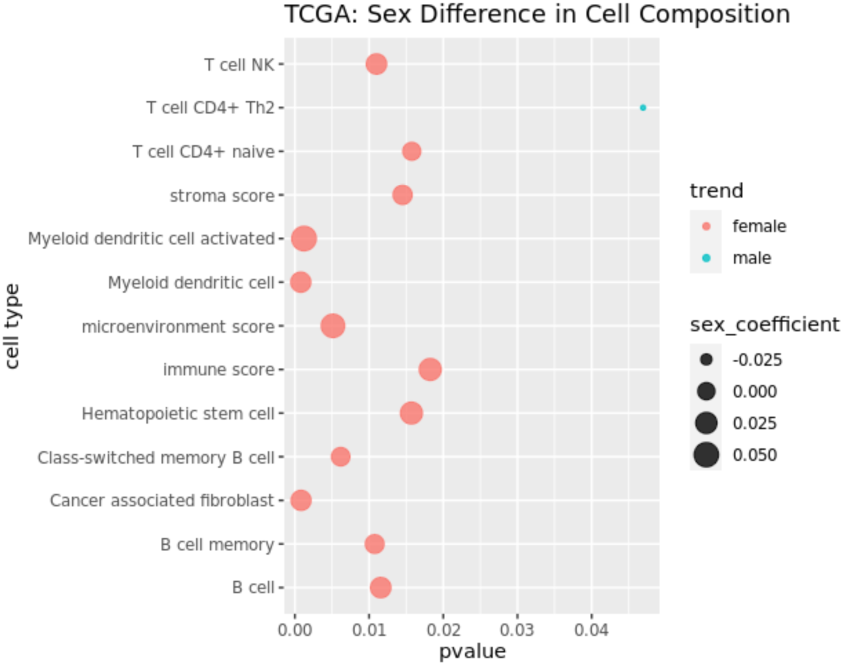
Sex Difference in immune and stromal cell composition in TCGA LUAD samples. Cell compositions are computed using “xcell”, which derives cell composition proportion of 36 immune and stromal, along with three composite scores: immune score, stroma score and microenvironment score. The bubbleplot shows only those cells that are significantly (p-value < 0.05) different in proportion in male and female tumor samples.

### Sex Difference in Chemotherapy

There is empirical evidence of significant sex differences in chemotherapy response [28] in LUAD, with females having better outcomes than males in most cases [6]. To explore this, we used networks only for patients who received chemotherapy and fit a Cox proportional hazard model to identify pathways with a sex-biased impact on survival. We find two pathways: drug metabolism CYP450 and metabolism of xenobiotics by CYP450. We observe that a higher targeting of both these pathways have a beneficial impact on prognosis in females receiving chemotherapy but for males receiving chemotherapy, we did not find any significant impact.

Within females, a higher targeting of two CYP450 pathways—drug metabolism (p-value 0.016) and metabolism of xenobiotics (p-value 0.052) was associated with better survival, while in males a differential targeting of these pathways did not have any impact on survival (p-value for metabolism of xenobiotics by CYP450 was 0.110 and p-value for drug metabolism CYP450 was 0.157). Previously, [20] found this same pattern of influence on the interaction between drug metabolism CYP450 targeting and chemotherapy treatment, in the context of colon cancer. Interestingly, these pathways did not have any significant impact on survival in treatment-naïve tumor samples, which indicates that gene regulatory network analysis has the power to predict the potential for individuals to respond to clinical interventions, including the use of chemotherapy agents.

### Sex Difference in Targeted Therapy

Cancer therapeutics targeting specific genes have also been observed to have a sex-biased impact on both dose-efficacy and dose-toxicity [55]. To understand how differential regulation of specific drug targets might contribute towards different efficacy of various cancer drugs in males and females with LUAD, we chose 28 genes commonly targeted by lung cancer drugs [29] for a closer analysis [**Figure 7**]. Among these genes, three showed significant (p-value less than 0.05) sex-bias in transcription factor targeting patterns: within nonsmokers *AKT2* showed higher targeting among females; *KRAS* and *IGF1R* showed higher targeting among males compared to females, irrespective of smoking status.

**Figure 7:**
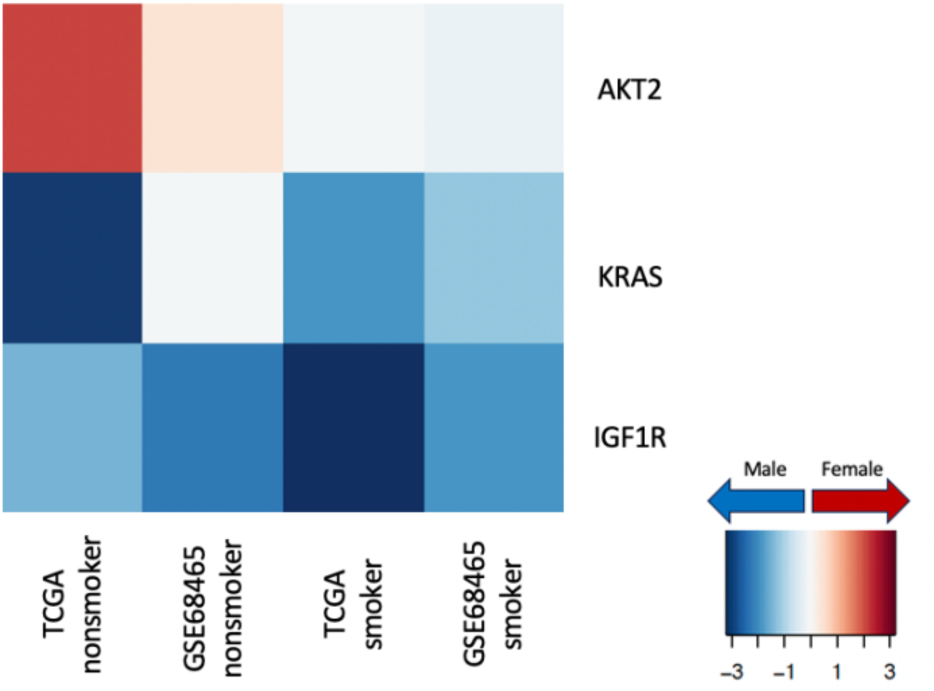
Sex difference in transcription factor targeting of genes commonly targeted by drugs in lung cancer in TCGA and validation data GSE68465, split by smoking status. The heatmap shows t-statistics corresponding to the sex coefficient from a limma analysis on the gene targeting score (indegree) (p-value < 0.05 for the sex coefficient). Genes with higher targeting in male samples are marked in blue and genes with higher targeting in female samples are marked in red.

Furthermore, to find potential targeted cancer therapeutics that might be more beneficial to individuals of one sex over the other, we used CLUEreg [30], a tool designed to match disease states to potentially therapeutic small molecule drugs based on differential regulation between tumor and healthy samples, and derived a list of small molecule drug candidates for both males and females. After cross-referencing these candidate drugs with the Genomics of Drug Sensitivity in Cancer (GDSC), we identified several small molecule drugs that might be beneficial for either males or females with LUAD. While several conventional cancer therapeutics such as Tanespimycin and Cisplatin appeared as potential drug candidates for both sexes, we found three drug candidates (Trametinib, Scriptaid/Vorinostat and Actinomycin-d/Dactinomycin) that had evidence of potential efficacy exclusively for females and one drug candidate (LBH-589/Panobinostat) exclusively for males; all four of these drugs are FDA approved.

Using GDSC dataset, we validated that female cell lines had higher sensitivity for Trametinib (p-value 0.00027 Mann-Whitney test), and male cell lines had higher sensitivity for Panobinostat (p-value 0.01396 Mann-Whitney test), as predicted by CLUEreg [**Figure 8**]. However, we did not find supporting evidence for sex differences in the efficacy of Vorinostat or Dactinomycin. This may be due to the relatively small number of cell lines of either sex that have been profiled and the innate variability among individuals in regulatory potential. Although preliminary, the validation of CLUEreg drug predictions using an independent cell line drug screening dataset confirms the value of using sex-specific changes of regulatory networks to identify therapeutics tailored to the patient sex.

**Figure 8:**
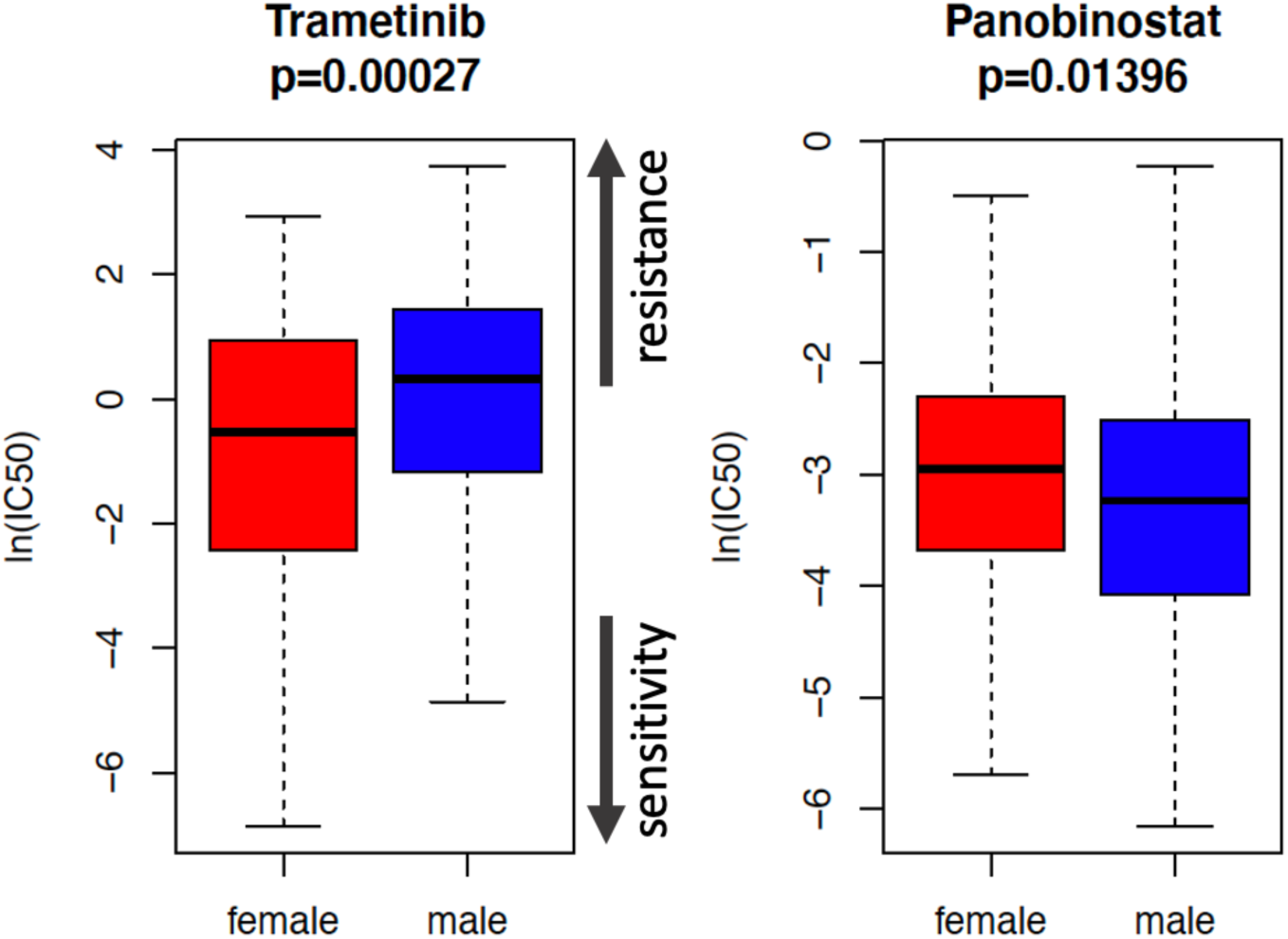
Validation of sex-specific therapeutics predicted by CLUEreg using GDSC drug sensitivity data. Boxplots of half maximal inhibitory concentration values (Log IC50) for male and female cell lines treated with Trametinib and Panobinostat, Mann-Whitney test.

## Discussion

LUAD, like many cancers, is known to differ between males and females in disease risk, development, progression, and response to therapy. While lifestyle differences, androgen and estrogen levels, and the genetic effects of different allosomes may play some role, the causes of these apparent sex differences remain largely unstudied. Although there are some differences in gene expression between males and females, both in healthy and tumor samples, these are largely confined to the sex chromosomes [20] and do not shed much light on mechanistic differences that might help explain the well-established clinical differences.

We applied methods to infer and compare gene regulatory network models to explore whether sex-specific regulatory patterns in healthy and LUAD samples might provide mechanistic explanations for sex-specific phenotypic differences in the disease. Using differential targeting analysis on individual-specific gene regulatory networks inferred using PANDA and LIONESS, we identified sex-bias in transcription factor targeting of biological pathways associated with cell proliferation, environmental carcinogen metabolism and immune response in healthy lungs, as well as in LUAD.

We found differences in regulatory processes controlling genes involved in cell proliferation and adhesion pathways, including many implicated in cancer, such as the hedgehog signaling pathway [31], WNT signaling pathway [32], notch signaling pathway [33] and ERBB signaling pathway [34]. Within healthy samples these pathways showed higher targeting in female nonsmokers and male smokers, whereas within tumor samples all these pathways were highly targeted in males, irrespective of smoking status. These differences in gene regulation may explain why females have a greater risk of developing LUAD, but the disease trajectory in males leads to more rapid progression and poorer outcomes.

Chemotherapy drugs such as carboplatin and paclitaxel has been observed to have sex difference in both efficacy and toxicity in non-small cell lung cancer, where females have more favorable prognosis than males [6]. Our analysis suggests that the differential response to chemotherapeutic agents might be associated to a differential targeting of drug metabolism CYP450 pathways. Among patients undergoing chemotherapy, we found that higher targeting of two CYP450 pathways, namely drug metabolism and xenobiotics metabolism, was associated to improved survival in females, while in males, differential targeting of these pathways did not have any significant impact on survival. A similar influence of drug metabolism CYP450 targeting on chemotherapy outcomes was previously identified in the context of colon cancer [20].

Not surprisingly, we also found sex-specific differences in the regulation of immune related processes, as well as proportion of infiltration of various immune cells within tumor samples. Not only do this shed light on cancer prognosis but might also elucidate towards a sex-biased response to various cancer immunotherapies [27], including PD1 and PDL1 inhibitors.

We identified that several genes for which targeted therapies exist, including *AKT2, IGF1R,* and *KRAS*, are differentially targeted between the sexes in our regulatory networks. While these genes have been extensively studied, there are virtually no published studies on potential sex differences in response to drugs targeting these genes. However, evidence for sex differences in response to targeted therapies is growing. It has been shown in a murine model that drugs targeting *IGF1R* (Insulin-like Growth Factor-1) improve lifespan with a reduction of neoplasm only in females [35], which aligns with our findings.

We identified four FDA-approved small-molecule drug candidates that might have a sex-biased efficacy: three drugs (Trametinib, Vorinostat and Dactinomycin) were identified exclusively for females and Panobinostat was identified exclusively for males. Using an independent database, we validated that female cell lines had indeed higher sensitivity for Trametinib, and male cell lines had higher sensitivity for Panobinostat. Trametinib targets *MAP2K1* [36], which showed higher targeting in males than females, based on our analysis of regulatory networks. Higher targeting of *MAP2K1* by transcription factors may reduce the effectiveness of cancer therapeutics targeting *MAP2K1* such as trametinib in males compared to females. Panobinostat is a histone deacetylase (HDAC) inhibitor [37]. HDAC inhibitors cause upregulation of the cell cycle gene *CDKN1A*, leading to cell cycle arrest [38, 39]. *CDKN1A* showed higher targeting by transcription factors in females than males. Higher targeting of *CDKN1A* by transcription factors may reduce the effectiveness of HDAC inhibitors such as Panobinostat in females compared to males. The validation of CLUEreg drug predictions using an independent cell line drug screening dataset underscores the potential of using gene regulatory networks to identify sex-specific cancer therapeutics.

It is essential to acknowledge that although during differential targeting analysis the data were adjusted for various clinical confounders such as age, race, smoking history, and clinical tumor stage, the analysis might still be influenced by cellular and genetic heterogeneity or unobserved clinical phenotypes and risk factors including the effect of hormones, lifestyle habits, environmental exposures, and family history. To establish causal conclusions regarding the effect of regulatory sex-differences in disease mechanism, further efforts are required to elucidate the relative contributions, as well as possible interactions between these factors and sex-biased gene regulatory patterns identified by our analysis.

In conclusion, our study highlights the substantial sex differences in gene regulatory patterns in healthy lung as well as LUAD. These distinctions not only bear relevance to disease susceptibility and prognosis but also hold promise for shaping sex-specific therapeutic responses and enhancing survival rates. Our findings emphasize the potential of harnessing sex-specific alterations in regulatory networks to develop personalized treatments and dosage protocols, tailored to each patient’s sex. Consequently, gene regulatory network inference emerges as a promising tool for designing sex-specific Precision Medicine approaches for LUAD as well as other diseases, to improve clinical outcomes for all individuals.

## Acknowledgements

This work was supported by grants from the National Institutes of Health: **ES, CMLR, MBG, VF, JF, KHS, PM, and JQ** were supported by R35CA220523; **MBG** and **JQ** were also supported by U24CA231846**; JQ** received additional support from P50CA127003; **JQ** and **DLD** were supported by R01HG011393; **KHS** and **DLD** were supported by R01HG125975 and P01HL114501; **KHS** was supported by T32HL007427; **CMLR** was supported by K01HL166376; **CMLR** and **ES** were also supported by the American Lung Association grant LCD-821824.

## Author Contributions

**ES:** Conceptualization, Data curation, Formal analysis, Investigation, Methodology, Software, Validation, Visualization, Writing – original draft; **MBG:** Conceptualization, Resources, Software, Writing – review & editing; **VF:** Conceptualization, Software, Writing – review & editing; **JF, KHS, PM:** Conceptualization, Writing – review & editing; **DLD:** Conceptualization, Funding acquisition, Supervision, Writing – review & editing; **JQ:** Conceptualization, Funding acquisition, Resources, Supervision, Writing – review & editing; **CMLR:** Conceptualization, Data curation, Funding acquisition, Methodology, Software, Supervision, Visualization, Writing – review & editing.

## STAR Methods

### Discovery Dataset

We downloaded uniformly processed RNA-Seq data from the Recount3 database [40] for two discovery datasets using the R package “recount3” (version 1.4.0) on May 26, 2022: (i) healthy lung tissue samples from the Genotype Tissue Expression (GTEx) Project [41] (version 8) and (ii) lung adenocarcinoma (LUAD) samples from The Cancer Genome Atlas (TCGA) [42]. Clinical data for GTEx samples were accessed from the dbGap website (https://dbgap.ncbi.nlm.nih.gov/) under study accession phs000424.v8.p2. Clinical data for TCGA samples were downloaded from Recount3. Throughout our analysis the GTEx samples will be referred to as “healthy lung samples.”

From 655 healthy lung samples in GTEx, we removed 77 samples because they were designated as “biological outliers” in the GTEx portal (https://gtexportal.org/) for various reasons (as described in https://gtexportal.org/home/faq). The remaining 578 samples (395 males, 183 females) were used in the analysis. We verified that the self-reported gender for GTEx samples aligned with the biological sex through a principal component analysis (PCA) of gene expression values of 36 genes on the Y chromosome [**Figure D.1**].

From the TCGA dataset, we removed two recurrent tumor samples and 59 samples from normal adjacent tissues, keeping only primary tumor samples. For individuals with multiple samples, we retained the sample with the highest sequencing depth. Finally, we also removed two samples annotated as “female” as these samples clustered with “male” samples using PCA for the Y chromosome as above [**Figure D.1**]. We also removed one sample with missing gender information. Subsequent analyses were performed on the remaining 513 primary lung adenocarcinoma tumor samples (238 males, 275 females).

We extracted TPM normalized gene expression data from both GTEx and TCGA using the “getTPM” function in the Bioconductor package “recount” (version 1.20.0) [43] in R (version 4.1.2). We excluded lowly expressed genes by removing those with counts <1 TPM in at least 10% of the samples in GTEx and TCGA combined, thus removing 36,360 annotated genes, and leaving 27,495 (including 36 Y genes and 884 X genes) genes for analysis. To build gene regulatory networks, we kept only genes that were present both in this filtered gene set and, in the TF-target gene regulatory prior used in PANDA and LIONESS (see section “differential targeting analysis using single-sample gene regulatory networks”). The remaining 27,189 genes, including genes on the sex chromosomes, were used for network inference and analysis. For female samples in both GTEx and TCGA, some genes on the Y chromosome have expression values due to mismapping of transcripts; we manually set Y chromosome gene expression values to “NA” for biological females in both data sets.

### Validation Dataset

We identified two independent studies from the Gene Expression Omnibus (GEO) for use in validating our findings: GSE47460 (hereafter referred to as LGRC) [44] and GSE68465 [45]. From the LGRC (downloaded on Feb 12, 2023) data, we used 108 samples (59 female and 49 male) annotated as “control” samples for validation. Gene expression data came from the Lung Genomics Research Consortium (LGRC) representing a subset of tissue samples from the Lung Tissue Research Consortium (LTRC) that showed no chronic lung disease by CT or pathology. This study used the Agilent-014850 Whole Human Genome Microarray 4×44K G4112F and Agilent-028004 SurePrint G3 Human GE 8×60K Microarray for gene expression profiling. Data from GSE68465 (downloaded on Jan 24, 2023) consisted of gene expression for lung adenocarcinoma primary tumor samples from 462 individuals. This study used Affymetrix Human Genome U133A Array for gene expression profiling. Nineteen samples were removed because of missing gender information. We also removed six samples annotated as “female” and five samples annotated as “male” based on PCA of expression of 65 Y genes [**Figure D.1**]. The remaining 432 samples (218 male and 214 female) were used in the final validation analysis.

Normalized expression data and clinical data were downloaded using the R package “GEOquery” version 2.62.2. For genes with multiple probe sets, we kept the probe with the highest standard deviation in expression across samples and the gene set was further filtered to remove any genes that did not overlap with those in the TF/target gene regulatory network prior. This left 13,575 genes in GSE47460 (LGRC) and 13,516 genes in GSE68465 that were used in subsequent analyses. The LGRC data did not show any batch effect and so no correction was used. The GSE68465 dataset contained LUAD specimens from the following sources: University of Michigan Cancer Center (100 samples), University of Minnesota VA/CALGB (77 samples), Moffitt Cancer Center (79 samples), Memorial Sloan-Kettering Cancer Center (104 samples), and Toronto/Dana-Farber Cancer Institute (82 samples). A principal component analysis on the gene expression data demonstrated distinct clusters corresponding to these sample source, thus exhibiting a strong batch effect; expression data was subsequently batch-corrected using the “ComBat” function implemented in the R package “sva” (version 3.42.0).

**Table 1** depicts the clinical characteristics of all the discovery and validation datasets.

**Table 1:**
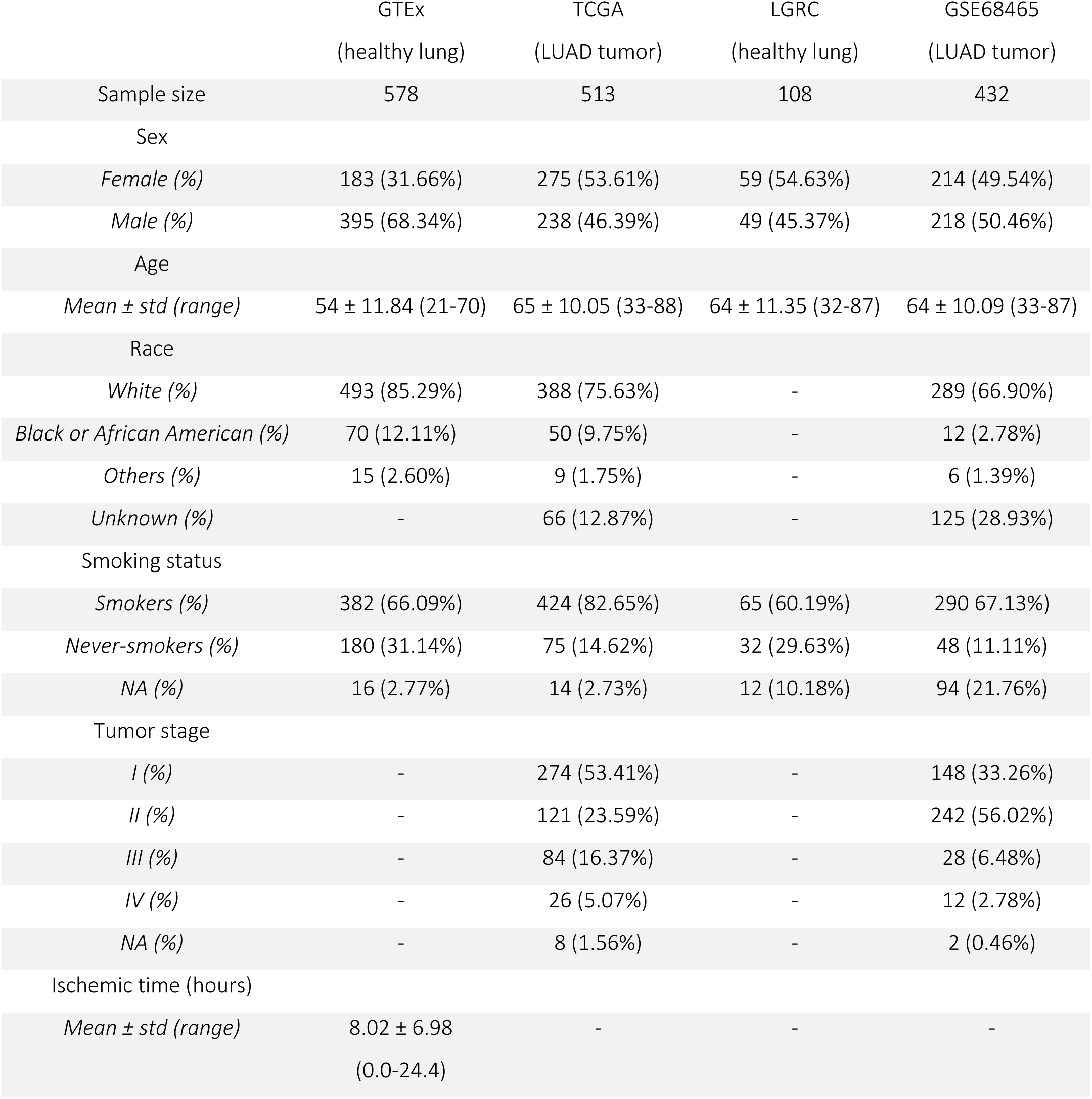
Clinical characteristics of the discovery and validation datasets. Clinical characteristics by sex are recorded in supplementary table S1.

|  | GTEx<br>(healthy lung) | TCGA<br>(LUAD tumor) | LGRC<br>(healthy lung) | GSE68465<br>(LUAD tumor) |
| --- | --- | --- | --- | --- |
| Sample size | 578 | 513 | 108 | 432 |
| Sex |  |  |  |  |
| *Female (%)* | 183 (31.66%) | 275 (53.61%) | 59 (54.63%) | 214 (49.54%) |
| *Male (%)* | 395 (68.34%) | 238 (46.39%) | 49 (45.37%) | 218 (50.46%) |
| Age |  |  |  |  |
| *Mean ± std (range)* | 54 ± 11.84 (21-70) | 65 ± 10.05 (33-88) | 64 ± 11.35 (32-87) | 64 ± 10.09 (33-87) |
| Race |  |  |  |  |
| *White (%)* | 493 (85.29%) | 388 (75.63%) | - | 289 (66.90%) |
| *Black or African American (%)* | 70 (12.11%) | 50 (9.75%) | - | 12 (2.78%) |
| *Others (%)* | 15 (2.60%) | 9 (1.75%) | - | 6 (1.39%) |
| *Unknown (%)* | - | 66 (12.87%) | - | 125 (28.93%) |
| Smoking status |  |  |  |  |
| *Smokers (%)* | 382 (66.09%) | 424 (82.65%) | 65 (60.19%) | 290 (67.13%) |
| *Never-smokers (%)* | 180 (31.14%) | 75 (14.62%) | 32 (29.63%) | 48 (11.11%) |
| *NA (%)* | 16 (2.77%) | 14 (2.73%) | 12 (10.18%) | 94 (21.76%) |
| Tumor stage |  |  |  |  |
| *I (%)* | - | 274 (53.41%) | - | 148 (33.26%) |
| *II (%)* | - | 121 (23.59%) | - | 242 (56.02%) |
| *III (%)* | - | 84 (16.37%) | - | 28 (6.48%) |
| *IV (%)* | - | 26 (5.07%) | - | 12 (2.78%) |
| *NA (%)* | - | 8 (1.56%) | - | 2 (0.46%) |
| Ischemic time (hours) |  |  |  |  |
| *Mean ± std (range)* | 8.02 ± 6.98<br>(0.0-24.4) | - | - | - |

### Differential Targeting Analysis using Single-sample Gene Regulatory Networks

We used PANDA [16] and LIONESS [17] to construct gene regulatory networks [**Figure 1**] for each sample in the discovery and validation datasets, using Python package netzooPy [46] version 0.9.10. In addition to the gene expression data obtained from the discovery and validation datasets, two other types of data were integrated to construct the networks: TF/target gene regulatory prior (derived by mapping TF motifs from the Catalog of Inferred Sequence Binding Preferences (CIS-BP) [47] to the promoter of their putative target genes) and protein-protein interaction data (using the interaction scores from StringDb v11.5 [48] between all TFs in the regulatory prior). Our TF/target gene regulatory prior consisted of 997 TFs targeting 61,485 ensemble gene IDs, corresponding to 39,618 unique gene symbols (HGNC), and the protein-protein interaction data contained the measure of interactions between these 997 TFs. We used sex-specific binary motif priors (1 representing the presence of a TF motif and 0 representing the absence of a TF motif on the promoter region of the gene) for males and females, where the male and female motifs were the same for autosomal and X chromosome genes, but motifs on the Y chromosome genes were set to 0 in the female prior. The procedure for deriving the motif prior and the PPI priors are given in the supplementary material. Regulatory networks were constructed for each of the discovery datasets and validation datasets separately for female and male samples. The final networks contained only genes overlapping between the TF/target gene motif prior and the corresponding gene expression dataset.

For each sample’s gene regulatory network, we computed the targeting score (or, in-degree) for each gene, which corresponds to the sum of incoming edge weights from all TFs to this gene. Gene targeting scores were compared between males and females using a linear regression model, while adjusting for relevant covariates: sex (Male and Female), race (White, Black or African American, Others and Unknown), age, smoking status (Ever-smoker and Never-smoker) and ischemic time for GTEx; sex (Male and Female), race (White, Black or African American, Others and Unknown), age, smoking status (Ever-smoker and Never-smoker) and tumor stage (stages I, II, III, IV and “NA”) for TCGA; using the R package limma (version 3.50.3) [49] and accounting for interaction between sex and smoking history (ever-smokers and never-smokers). In the LGRC dataset we adjusted for age and smoking status and in GSE68465 we adjusted for age, race, tumor stage and smoking status, while simultaneously considering interaction between sex and smoking history (ever-smokers and never-smokers) for each validation dataset.

### Pathway Enrichment Analysis

A gene set enrichment analysis was performed separately for individuals with different smoking histories using the ranked t-statistics of the coefficient for sex derived from the limma analysis (**Figure 1**). We used pre-ranked Gene Set Enrichment Analysis (GSEA) in the R package “fgsea” (version 1.20.0) [50] and gene sets from the Kyoto Encyclopedia of Genes and Genomes (KEGG) pathway database [51] (“c2.cp.kegg.v2022.1.Hs.symbols.gmt”), downloaded from the Molecular Signatures Database (MSigDB) (http://www.broadinstitute.org/gsea/msigdb/collections.jsp). Only gene sets of sizes greater than 15 and less than 500 were considered, after filtering out genes which are not present in the expression dataset, which limited our analysis to 176 gene sets. Multiple testing corrections were performed using the Benjamini-Hochberg procedure [52].

### Survival Analysis

For each biological pathway, the pathway targeting score was computed as the mean indegree of all genes in the pathway. For survival analysis we used the R package “survival” (version 3.2.13) and fit Cox proportional hazard model (“coxph”) for the TCGA data to investigate the effect of transcription factor targeting of different KEGG pathways on survival outcome, while adjusting for age, sex, race, smoking status, tumor stage, and chemotherapy status (yes, no and “NA”).

### Immune Infiltration Analysis

We used “xcell” [53] on the TPM-normalized GTEx and TCGA gene expression data with R package “immunedeconv” (version 2.1.0) to infer immune and stromal cell composition in tumor samples. For every cell type, to quantify whether cell type proportion in tumor are variable by sex, we fit a linear model to predict cell type proportion by sex, while adjusting for age, race, smoking status, and clinical tumor stage.

### Finding Small Molecule Drugs with CLUEreg

We identified genes that are differentially targeted between tumor and healthy samples, using linear models on gene targeting scores from GTEx and TCGA data through R package “limma”. We accounted for the interaction between sex and disease status (tumor versus healthy), while adjusting for clinical covariates that were available for both GTEx and TCGA, i.e., sex, age, race, and smoking status. Genes were ranked by the adjusted p-values (smallest to largest) from the limma analysis and all genes significantly differentially targeted (at FDR cutoff 0.05) were chosen for males and females separately. The selected differentially targeted genes were split between “high” and “low” targeted based on whether they were more highly targeted in tumor (high) samples or in healthy (low) samples and subsequently used as input to CLUEreg [30] (https://grand.networkmedicine.org/), a web application designed to match disease states to potential small molecule therapeutics, based on the characteristics of the regulatory networks. CLUEreg produced a list of 100 small molecule drug candidates most suitable for reversing the gene targeting patterns in tumor to resemble the gene targeting patterns in healthy samples.

To validate CLUEreg predictions, we used gene expression and drug response data from cancer cell lines in the Genomics of Drug Sensitivity in Cancer (GDSC) [54] dataset, removing cell lines from reproductive cancer types. We classified cell lines as male (n=227) or female (n=264) groups considering both expression of the Y chromosome genes (gene expression data from GDSC) and the reported gender of the individual from whom the cell line was derived (Sanger Cell Model Passports, https://cellmodelpassports.sanger.ac.uk/downloads). To test whether drug sensitivity varies by sex, we combined technical replicates by median of log IC50 and compared the log IC50 values reported by GDSC (half maximal inhibitory concentration) between male and female cell lines using Wilcoxon-Mann-Whitney test.

## Resource Availability

### Lead Contact

Further information and requests for resources should be directed to and will be fulfilled by the lead contact Camila M. Lopes-Ramos

### Materials Availability

This study did not generate new unique reagents.

## Data and Code Availability

Raw data to construct gene regulatory networks and other analysis were downloaded from open-source databases dbGap, Recount3, GEO, STRINGdb, CIS-BP and GDSC. Processed data are available upon request.

Sample-specific gene regulatory networks will be available in the GRAND database (https://grand.networkmedicine.org) upon acceptance.

R codes for all downstream analysis are available on a GitHub public repository: https://github.com/Enakshi-Saha/Sex-Differences-Lung-Adenocarcinoma

A notebook describing differential targeting analysis on the TCGA data will be available on Netbooks [55]: http://netbooks.networkmedicine.org upon acceptance.

## Supplemental information

A. **Designing Sex-specific Transcription Factor-Gene Motif Prior**
B. **Designing Protein-protein Interaction Prior**
C. **Sex Difference in anti PD-1 and anti PDL-1 Inhibitors in Non-small Cell Lung Cancer**
D. **Additional Figures**

**Figure D.1: Defining biological sex based on sex chromosome complement.** Scatterplot of first two principal components of Y chromosome gene expression in GTEx (top left), TCGA (top right), LGRC (bottom left) and GSE68465 (bottom right).

**Figure D.2:** Sex difference in LGRC control lung samples within nonsmokers and smokers. Normalized enrichment scores (NES) from GSEA using KEGG pathways are shown for all pathways that have significant (adjusted p-value < 0.05) sex difference among either nonsmokers or smokers in LGRC. Pathways with higher targeting in male are marked blue and pathways with higher targeting in female are marked red. Green boxes highlight pathways associated with cell proliferation and brown boxes highlight pathways associated with environmental carcinogen metabolism.

**Figure D.3**: Sex difference in tumor samples from the validation data GSE68465 within nonsmokers and smokers. Normalized enrichment scores (NES) from GSEA using KEGG pathways are shown for all pathways that have significant (adjusted p-value < 0.05) sex difference among either nonsmokers or smokers (in TCGA). Pathways with higher targeting in male are marked blue and pathways with higher targeting in female are marked red. Green boxes highlight pathways associated with cell proliferation and purple boxes highlight pathways associated with immune response.

**Figure D.4**: Sex difference in immune and stromal cell composition in GTEx samples: nonsmokers (left) and smokers (right). Cell compositions are computed using “xcell”, which derives cell composition proportion of 36 immune and stromal, along with three composite scores: immune score, stroma score and microenvironment score. The bubbleplot shows only those cells that are significantly (p-value < 0.05) different in proportion in male and female samples.

**E. Additional Tables**

**Table E.1:** Distribution of Clinical Variables by Sex in GTEx.

**Table E.2:** Distribution of Clinical Variables by Sex in TCGA.

**Table E.3:** Distribution of Clinical Variables by Sex in LGRC.

**Table E.4:** Distribution of Clinical Variables by Sex in GSE68465.

---

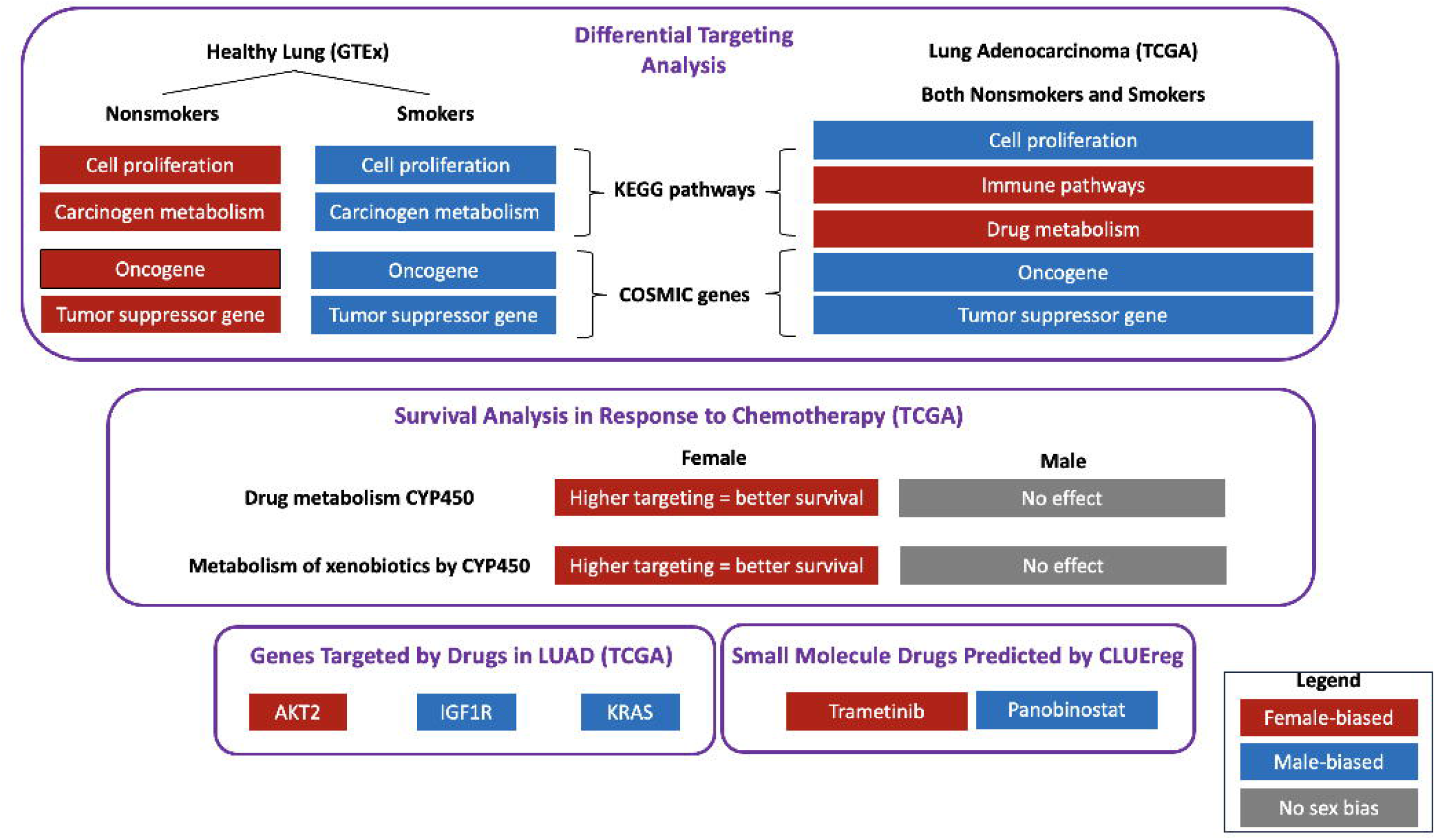

## Bibliography

[1] A. Jemal, K. D. Miller, J. Ma, R. L. Siegel, S. A. Fedewa, F. Islami, S. S. Devesa and M. J. Thun, "Higher lung cancer incidence in young women than young men in the United States," New England Journal of Medicine, vol. 378, no. 21, pp. 1999–2009, 2018.

[2] J. Hellyer and M. Patel, "Sex disparities in lung cancer incidence: validation of a long-observed trend," Transl Lung Cancer Res., vol. 8, no. 4, pp. 543–545, 2019 Aug.

[3] M. M. Fidler-Benaoudia, L. A. Torre, F. Bray, J. Ferlay and A. Jemal, "Lung cancer incidence in young women vs. young men: a systematic analysis in 40 countries," International journal of cancer, vol. 147, no. 3, pp. 811–819, 2020.

[4] J. M. Siegfried, "Women and lung cancer: does oestrogen play a role?," The lancet oncology, vol. 2, no. 8, pp. 506–513, 2001.

[5] N. Mederos, A. Friedlaender, S. Peters and A. Addeo, "Gender-specific aspects of epidemiology, molecular genetics and outcome: lung cancer," ESMO Open, vol. 5, p. e000796, 2020 Nov.

[6] H. Yamamoto, I. Sekine, K. Yamada, H. Nokihara, N. Yamamoto, H. Kunitoh, Y. Ohe and T. Tamura, "Gender differences in treatment outcomes among patients with non-small cell lung cancer given a combination of carboplatin and paclitaxel," Oncology., vol. 75, no. 3-4, pp. 169–74, 2008.

[7] F. Conforti, L. Pala, V. Bagnardi, G. Viale, T. De Pas, E. Pagan and E. Pennacchioli et al., "Sex-based heterogeneity in response to lung cancer immunotherapy: a systematic review and meta-analysis," JNCI: Journal of the National Cancer Institute, vol. 111, no. 8, pp. 772–781, 2019.

[8] C. M. Dresler, C. Fratelli, J. Babb, L. Everley, A. A. Evans and M. L. Clapper, "Gender differences in genetic susceptibility for lung cancer," Lung cancer, vol. 30, no. 3, pp. 153–160, 2000.

[9] Y. Omoto, Y. Kobayashi and K. Nishida et al, "Expression, function, and clinical implications of the estrogen receptor beta in human lung cancers," Biochemical and biophysical research communications, vol. 285, no. 2, pp. 340–347, 2001.

[10] S. Mollerup, K. Jorgensen and G. Berge et al, "Expression of estrogen receptors alpha and beta in human lung tissue and cell lines," Lung Cancer, vol. 37, no. 2, pp. 153–159, 2002.

[11] C. Stapelfeld, C. Dammann and E. Maser, "Sex-specificity in lung cancer risk," International journal of cancer, vol. 146, no. 9, pp. 2376-2382, 2020.

[12] T. McLemore, S. Adelberg and M. Liu et al, "Expression of CYP1A1 gene in patients with lung cancer: Evidence for cigarette smoke-induced gene expression in normal lung tissue and for altered gene regulation in primary pulmonary carcinomas," JNCI: Journal of the National Cancer Institute, vol. 82, no. 16, pp. 1333–1339, 1990.

[13] J. Patel, P. Bach and M. Kris, "Lung cancer in US women: A contemporary epidemic.," JAMA, vol. 291, no. 14, pp. 1763–1768, 2004.

[14] X. Li, S. Wei, L. Deng and H. Tao et. al, "Sex-biased molecular differences in lung adenocarcinoma are ethnic and smoking specific," BMC Pulmonary Medicine, vol. 23, no. 1, pp. 1–14, 2023.

[15] Y. Yuan, L. Liu, H. Chen and Y. Wang et. al, "Comprehensive characterization of molecular differences in cancer between male and female patients," Cancer cell, vol. 29, no. 5, pp. 711–722, 2016.

[16] K. Glass, C. Huttenhower, J. Quackenbush and G.-C. Yuan, "Passing messages between biological networks to refine predicted interactions," PloS one, vol. 8, no. 5, p. e64832, 2013.

[17] M. L. Kuijjer, M. G. Tung, G. Yuan, J. Quackenbush and K. Glass, "Estimating sample-specific regulatory networks," Iscience, vol. 14, pp. 226–240, 2019.

[18] C. Lopes-Ramos, J. Quackenbush and D. DeMeo, "Genome-Wide Sex and Gender Differences in Cancer," Front Oncol., vol. 23, no. 10, p. 597788, 2020 Nov.

[19] K. Glass, J. Quackenbush, E. Silverman, B. Celli, S. Rennard and G.-C. Yuan et al, "Sexuallydimorphic targeting of functionally-related genes in COPD," BMC Syst Biol., vol. 8, pp. 1–17, 2014.

[20] C. M. Lopes-Ramos, M. L. Kuijjer, S. Ogino, C. S. Fuchs, D. L. DeMeo, K. Glass and J. Quackenbush, "Gene regulatory network analysis identifies sex-linked differences in colon cancer drug metabolism," Cancer research, vol. 78, no. 19, pp. 5538–5547, 2018.

[21] C. M. Lopes-Ramos, C.-Y. Chen and M. L. Kuijjer et. al, "Sex differences in gene expression and regulatory networks across 29 human tissues," Cell reports, vol. 31, no. 12, 2020.

[22] S. Bamford, E. Dawson, S. Forbes, J. Clements, R. Pettett, A. Dogan, A. Flanagan, J. Teague, P. A. Futreal, M. R. Stratton and others, "The COSMIC (Catalogue of Somatic Mutations in Cancer) database and website," British journal of cancer, vol. 91, no. 2, pp. 355--358, 2004.

[23] D. Cheng, J. Li and S. Li, "CNOT 1 cooperates with LMNA to aggravate osteosarcoma tumorigenesis through the Hedgehog signaling pathway," Molecular Oncology, vol. 11, no. 4, pp. 388–404, 2017.

[24] J. Lee, J. H. Jung and J. Hwang et. al, "CNOT2 is critically involved in atorvastatin induced apoptotic and autophagic cell death in non-small cell lung cancers," Cancers, vol. 11, no. 10, p. 1470, 2019.

[25] Y.-T. Shirai, A. Mizutani and S. Nishijima, "CNOT3 targets negative cell cycle regulators in non-small cell lung cancer development," Oncogene, vol. 38, no. 14, pp. 2580–2594, 2019.

[26] Y. Zhang and H. Ma, "LncRNA HOXD-AS2 regulates miR-3681-5p/DCP1A axis to promote the progression of non-small cell lung cancer," ournal of Thoracic Disease, vol. 15, no. 3, p. 1289, 2023.

[27] A. Grassadonia, I. Sperduti and P. Vici et. al, "Effect of Gender on the Outcome of Patients Receiving Immune Checkpoint Inhibitors for Advanced Cancer: A Systematic Review and Meta-Analysis of Phase III Randomized Clinical Trials," J Clin Med., vol. 7, no. 12, p. 542, 2018 Dec 12.

[28] A. Wagner, "Sex differences in cancer chemotherapy effects, and why we need to reconsider BSA-based dosing of chemotherapy," ESMO Open, vol. 5, no. 5, 2020 Sep.

[29] E. Shtivelman, T. Hensing, G. R. Simon, P. A. Dennis, G. A. Otterson, R. Bueno and R. Salgia, "Molecular pathways and therapeutic targets in lung cancer," Oncotarget, vol. 5, no. 6, p. 1392, 2014.

[30] M. Ben Guebila, C. M. Lopes-Ramos and D. Weighill et. al, "GRAND: a database of gene regulatory network models across human conditions," Nucleic Acids Research, vol. 50, no. D1, pp. D610–D621, 2022.

[31] E. Giroux-Leprieur, A. Costantini, V. W. Ding and B. He, "Hedgehog signaling in lung cancer: From oncogenesis to cancer treatment resistance," International Journal of Molecular Sciences, vol. 19, no. 9, p. 2835, 2018.

[32] D. J. Stewart, "Wnt signaling pathway in non–small cell lung cancer," JNCI: Journal of the National Cancer Institute, vol. 106, no. 1, 2014.

[33] B. Zou, X. Zhou, S. Lai and J. Liu, "Notch signaling and non-small cell lung cancer," Oncology letters, vol. 15, no. 3, pp. 3415-3421, 2018.

[34] M. O. Hoque, M. Brait, E. Rosenbaum, M. L. Poeta, P. Pal, S. Begum and S. Dasgupta et al., "Genetic and epigenetic analysis of erbB signaling pathway genes in lung cancer," Journal of Thoracic Oncology, vol. 5, no. 12, pp. 1887–1893, 2010.

[35] K. Mao, G. F. Quipildor, T. Tabrizian, A. Novaj, F. Guan, R. O. Walters, F. Delahaye, G. B. Hubbard, Y. Ikeno, K. Ejima and others, "Late-life targeting of the IGF-1 receptor improves healthspan and lifespan in female mice," Nature communications, vol. 9, no. 1, p. 2394, 2018.

[36] C. J. Wright and P. L. McCormack, "Trametinib: first global approval," Drugs, vol. 73, pp. 1245–1254, 2013.

[37] N. R. Srinivas, "Clinical pharmacokinetics of panobinostat, a novel histone deacetylase (HDAC) inhibitor: review and perspectives," Xenobiotica, vol. 47, no. 4, pp. 354–368, 2017.

[38] A. A. Mensah and I. Kwee et. al, "Novel HDAC inhibitors exhibit pre-clinical efficacy in lymphoma models and point to the importance of CDKN1A expression levels in mediating their anti-tumor response," Oncotarget, vol. 7, no. 2015, p. 5059, 6.

[39] V. M. Richon and T. W. Sandhoff et. al, "Histone deacetylase inhibitor selectively induces p21WAF1 expression and gene-associated histone acetylation," Proceedings of the National Academy of Sciences, vol. 97, no. 18, pp. 10014–10019, 2000.

[40] C. Wilks, S. Zheng and F. Chen, et al., "recount3: summaries and queries for large-scale RNA-seq expression and splicing," Genome Biol, vol. 22, p. 323, 2021.

[41] J. Lonsdale, J. Thomas, M. Salvatore, R. Phillips and E. Lo et al., "The genotype-tissue expression (GTEx) project," Nature genetics, vol. 45, no. 6, pp. 580--585, 2013.

[42] The Cancer Genome Atlas Research Network. Weinstein, J., Collisson, E. et al., "The cancer genome atlas pan-cancer analysis project," Nat Genet, vol. 45, pp. 1113-1120, 2013.

[43] L. Collado-Torres, A. Nellore and K. Kammers et al., "Reproducible RNA-seq analysis using recount2," Nat Biotechnol, vol. 35, p. 319–321, 2017.

[44] S. Kim, J. Herazo-Maya and D. Kang et al., "Integrative phenotyping framework (iPF): integrative clustering of multiple omics data identifies novel lung disease subphenotypes," BMC Genomics, vol. 16, p. 924, 2015.

[45] Director’s Challenge Consortium for the Molecular Classification of Lung Adenocarcinoma, "Gene expression–based survival prediction in lung adenocarcinoma: a multi-site, blinded validation study," Nat Med, vol. 14, p. 822–827, 2008.

[46] M. Ben Guebila, T. Wang, C. M. Lopes-Ramos, V. Fanfani, D. Weighill, R. Burkholz and D. Schlauch et al., "The Network Zoo: a multilingual package for the inference and analysis of gene regulatory networks," Genome Biology, vol. 24, no. 1, pp. 1–23, 2023.

[47] M. Weirauch, A. Yang, M. Albu, A. Cote, A. Montenegro-Montero and P. Drewe et al, "Determination and inference of eukaryotic transcription factor sequence specificity," Cell., vol. 158, pp. 1431–43, 2014.

[48] D. Szklarczyk, A. L. Gable, K. C. Nastou and D. Lyon et. al, "The STRING database in 2021: customizable protein-protein networks, and functional characterization of user-uploaded gene/measurement sets," Nucleic Acids Research (Database issue), vol. 49, 2021.

[49] "limma powers differential expression analyses for RNA-sequencing and microarray studies," Nucleic Acids Res, vol. 43, p. e47, 2015.

[50] G. Korotkevich, V. Sukhov, N. Budin, B. Shpak, M. N. Artyomov and A. Sergushichev., "Fast gene set enrichment analysis," BioRxiv, p. 060012, 2016.

[51] M. Kanehisa, "The KEGG database," *‘*In Silico’Simulation of Biological Processes: Novartis Foundation Symposium 247, vol. 247, pp. 91–103, 2002.

52. Y. Benjamini and Hochberg, Y, "Controlling the False Discovery Rate: A Practical and Powerful Approach to Multiple Testing," J R Stat Soc Ser B. Blackwell Publishing for the Royal Statistical Society, vol. 57, p. 289–300, 1995.

[53] D. Aran, Z. Hu and A. Butte, "xCell: digitally portraying the tissue cellular heterogeneity landscape," Genome Biol, vol. 18, p. 220, 2017.

[54] W. Yang, J. Soares, P. Greninger and E. J. Edelman et. al, "Genomics of Drug Sensitivity in Cancer (GDSC): a resource for therapeutic biomarker discovery in cancer cells," Nucleic acids research, vol. 41, no. D1, pp. D955–D961, 2012.

[55] M. Ben Guebila, D. Weighill and C. M. Lopes-Ramos et. al, "An online notebook resource for reproducible inference, analysis and publication of gene regulatory networks," Nature methods, vol. 19, no. 5, pp. 511–513, 2022.

[56] C. E. Grant, T. L. Bailey and W. S. Noble, "FIMO: scanning for occurrences of a given motif,," Bioinformatics, vol. 27, no. 7, 2011.

[57] R. Herbst, P. Baas and D. W. Kim et. al, "Pembrolizumab versus docetaxel for previously treated, PD-L1-positive, advanced non-small-cell lung cancer (KEYNOTE-010): A randomised controlled trial," Lancet, vol. 387, p. 1540–1550, 2016.

[58] M. Reck, D. Rodríguez-Abreu and A. Robinson et. al, "Pembrolizumab versus chemotherapy for PD-L1-positive non-small-cell lung cancer," N. Engl. J. Med., vol. 375, pp. 1823–1833, 2016.

[59] A. Rittmeyer, F. Barlesi, D. Waterkamp and K. Park et. al, "Atezolizumab versus docetaxel in patients with previously treated non-small-cell lung cancer (OAK): A phase 3, open-label, multicentre randomised controlled trial," Lancet, vol. 389, p. 255–265, 2017.

[60] D. Carbone, M. Reck, L. Paz-Ares, B. Creelan and L. Horn et. al, "First-line nivolumab in stage iv or recurrent non-small-cell lung cancer," N. Engl. J. Med., vol. 376, p. 2415–2426, 2017.

[61] S. Antonia, A. Villegas and D. Daniel et. al, "Durvalumab after chemoradiotherapy in stage III non-small-cell lung cancer," N. Engl. J.Med., vol. 377, p. 1919–1929, 2017.

[62] X. Sun, T. Zhang, M. Li, L. Yin and J. Xue, "Immunosuppressive B cells expressing PD-1/PD-L1 in solid tumors: A mini review," QJM. 2019 Jun 26, p. hcz162, 2019 Jun 26.

[63] F. Mueller, B. Büchel and D. Köberle et. al, "Gender-Specific elimination of continuous-infusional 5-fluorouracil in patients with gastrointestinal malignancies: results from a prospective population pharmacokinetic study," Cancer Chemother Pharmacol, 2013.

[64] Z. Zhang, L. Xu, L. Huang, T. Li, J. Y. Wang, C. Ma, X. Bian, X. Ren, H. Li and X. Wang, "Glutathione S-Transferase Alpha 4 Promotes Proliferation and Chemoresistance in Colorectal Cancer Cells," Frontiers in Oncology, vol. 12, 2022.

[65] W. Li, W. Yue and L. Zhang et al., "Polymorphisms in GSTM1, CYP1A1, CYP2E1, and CYP2D6 are Associated with Susceptibility and Chemotherapy Response in Non-small-cell Lung Cancer Patients," Lung, vol. 190, pp. 91–98, 2012.

[66] C. Rodriguez-Antona and M. Ingelman-Sundberg, "Cytochrome P450 pharmacogenetics and cancer," Oncogene, vol. 25, no. 11, pp. 1679–91, 2006 Mar.

[67] A. K. Croker and A. L. Allan, "Inhibition of aldehyde dehydrogenase (ALDH) activity reduces chemotherapy and radiation resistance of stem-like ALDH^hi CD44+ human breast cancer cells," Breast Cancer Res Treat, vol. 133, p. 75–87, 2012.

[68] S. Duan, B. Gong, P. Wang, H. Huang, L. Luo and F. Liu, "Novel prognostic biomarkers of gastric cancer based on gene expression microarray: COL12A1, GSTA3, FGA and FGG," Molecular medicine reports, vol. 18, no. 4, pp. 3727–3736, 2018.

[69] L. Peng, L. Zhuang, K. Lin, Y. Yao, Y. Zhang, T. Arumugam and T. Fujii et. al, "Downregulation of GSTM2 enhances gemcitabine chemosensitivity of pancreatic cancer in vitro and in vivo," Pancreatology, vol. 21, no. 1, pp. 115–123, 2021.

[70] C. Tibaldi, I. Stasi and E. Baldini, "Oncogene-addicted non-small-cell lung cancer in women: a narrative review of the importance of gender-related differences in treatment outcome," 2022.

[71] A. D. Wagner, S. Oertelt-Prigione and A. Adjei et. al, "Gender medicine and oncology: report and consensus of an ESMO workshop," Annals of Oncology, vol. 30, no. 12, pp. 1914–1924, 2019.

[72] M. Scheffler, A. Holzem and A. Kron et. al, "Co-occurrence of targetable mutations in Non-small cell lung cancer (NSCLC) patients harboring MAP2K1 mutations," Lung Cancer, vol. 144, pp. 40–48, 2020.

