## Supplementary Material for "Gene regulatory Networks Reveal Sex Difference in Lung Adenocarcinoma"

### Supplementary Materials

Enakshi Saha[1], Marouen Ben Guebila[1], Viola Fanfani[1], Jonas Fischer[1], Katherine Hoff-Shutta[1,2], Panagiotis Mandros[1], Dawn L DeMeo[2,3], John Quackenbush[1,2,4], Camila M Lopes-Ramos[1,2,3]

[1] Department of Biostatistics, Harvard T. H. Chan School of Public Health, Boston, MA 02115, USA

[2] Channing Division of Network Medicine, Brigham and Women's Hospital, Boston, MA, USA 02115

[3] Department of Medicine, Harvard Medical School, Boston, MA 02115, USA

[4] Department of Data Science, Dana-Farber Cancer Institute, Boston, MA 02115, USA

**Corresponding Author:** Camila M Lopes-Ramos

### A. Designing Sex-specific Transcription Factor-Gene Motif Prior

The prior regulatory network is a binary network of transcription factors to their target genes, where the edges (0 or 1) indicate whether a transcription factor motif exists in a target gene's promoter. To create the prior regulatory network, we downloaded Homo sapiens transcription factor motifs with direct/inferred evidence from the Catalog of Inferred Sequence Binding Preferences CIS-BP Build 2.0 (http://cisbp.ccbr.utoronto.ca). We mapped these transcription factor position weight matrices (PWM) to the human genome (hg38) using FIMO [56] and retained highly significant matches (p<10-5) that occurred within the promoter regions of Ensembl genes (Gencode v39; annotations downloaded from http://genome.ucsc.edu/cgi-bin/hgTables); promoter regions were defined as [-750; +250] base pairs around the transcription start site (TSS). This process resulted in an initial map of potential regulatory interactions involving 997 transcription factors targeting 61,485 genes. To statistically compare networks, the same set of edge combinations need to be included in both sexes, therefore we created sex-informed transcription factor regulatory priors to account for the lack of Y chromosome genes in females. In the female regulatory prior, edges from or to Y chromosome genes were downweighed to zero, which consisted of 52,266 edges.

### B. Designing Protein-protein Interaction Prior

We used the STRINGdb Bioconductor package [48] to access and download PPI data from the StringDB database (STRING.version 11.5). We filtered the PPI data to keep only those interactions present between transcription factors in the prior network (score threshold index of 0). PPI scores were normalized by dividing them by 1000 to have a uniform range between 0 and 1 for the PPI and the motif prior network. We set transcription factor self-interaction equal to one. Since PPI are undirected, we converted the data into a symmetric form.

### C. Sex Difference in anti PD-1 and anti PDL-1 Inhibitors in Non-small Cell Lung Cancer

**Table C.1:** Sex difference in response to cancer therapeutics targeting PD-1 and PDL-1. All studies are based on non-small cell lung cancer clinical trials. The last column reports the hazard ratio for overall survival or progression-free survival. The following information is obtained from [27], except the target information is extracted from the corresponding clinical trial.

| Clinical Trial | Target | Treatment | No. Samples | | Hazard Ratio |
|---|---|---|---|---|---|
| | | | Male | Female | |
| [57] | PD-1 | Pembrolizumab | 425 | 266 | Male: 0.7 |
| | | Docetaxel | 209 | 134 | Female: 1.02 |
| [58] | PD-1 | Pembrolizumab | 92 | 62 | Male: 0.39 |
| | | Chemotherapy | 95 | 56 | Female: 0.75 |
| [59] | PDL-1 | Atezolizumab | 261 | 164 | Male: 0.79* |
| | | Docetaxel | 259 | 166 | Female: 0.64* |
| [60] | PD-1 | Nivolumb | 184 | 87 | Male: 1.05 |
| | | Chemotherapy | 148 | 122 | Female: 1.36 |
| [61] | PDL-1 | Durvalumab | 334 | 142 | Male: 0.54 |
| | | Placebo | 166 | 71 | Female: 0.54 |

*Hazard ratio is computed based on overall survival, for all other studies hazard ratio is computed based on progression free survival.

### D. Additional Figures

**Figure D.1: Defining biological sex based on sex chromosome complement.** Scatterplot of first two principal components of Y chromosome gene expression in GTEx (top left), TCGA (top right), LGRC (bottom left) and GSE68465 (bottom right).

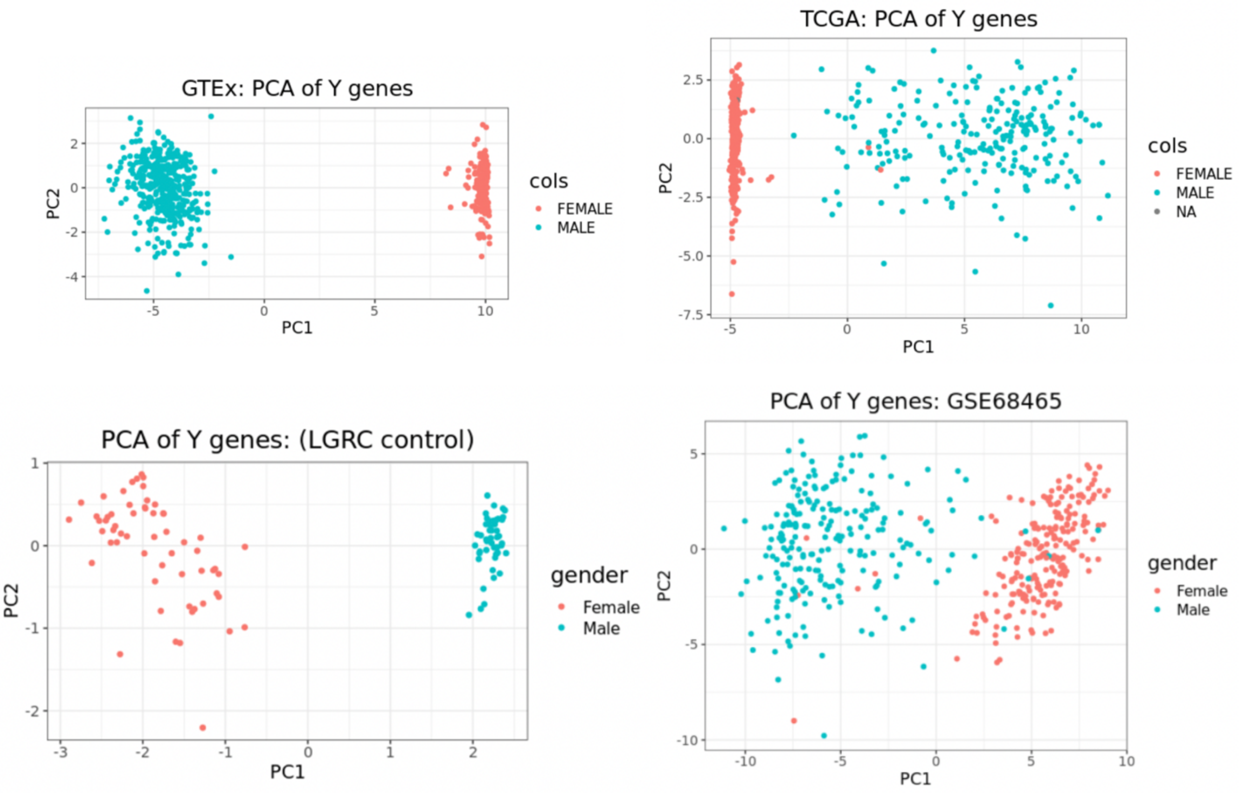
GTEx: PCA of Y genes
PC1
PC2
cols
FEMALE
MALE
TCGA: PCA of Y genes
PC1
PC2
cols
FEMALE
MALE
NA
PCA of Y genes: (LGRC control)
PC1
PC2
gender
Female
Male
PCA of Y genes: GSE68465
PC1
PC2
gender
Female
Male

**Figure D.2:** Sex difference in LGRC control lung samples within nonsmokers and smokers. Normalized enrichment scores (NES) from GSEA using KEGG pathways are shown for all pathways that have significant (adjusted p-value < 0.05) sex difference among either nonsmokers or smokers in LGRC. Pathways with higher targeting in male are marked blue and pathways with higher targeting in female are marked red. Green boxes highlight pathways associated with cell proliferation and brown boxes highlight pathways associated with environmental carcinogen metabolism.

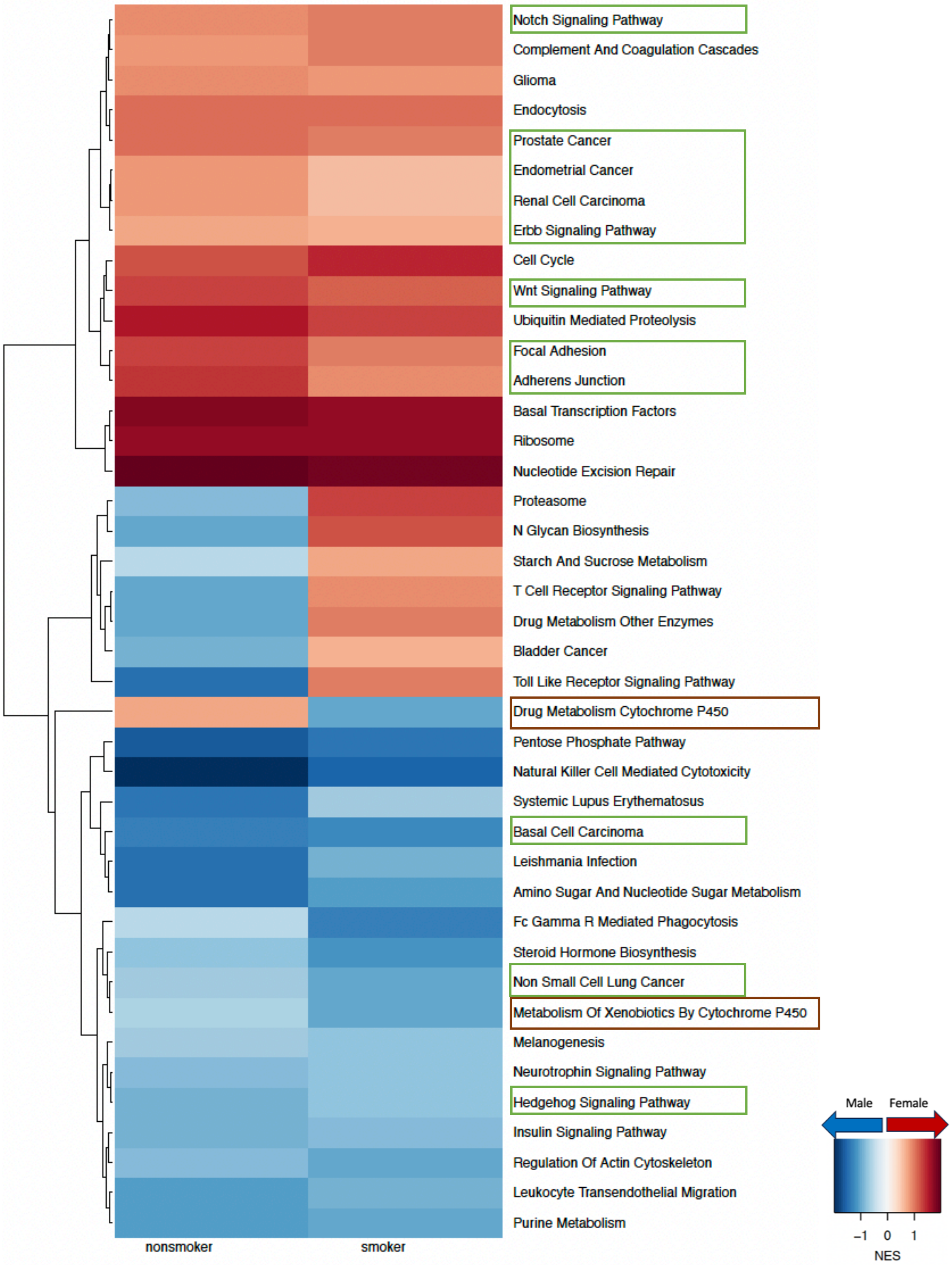

**Figure D.3**: Sex difference in tumor samples from the validation data GSE68465 within nonsmokers and smokers. Normalized enrichment scores (NES) from GSEA using KEGG pathways are shown for all pathways that have significant (adjusted p-value < 0.05) sex difference among either nonsmokers or smokers (in TCGA). Pathways with higher targeting in male are marked blue and pathways with higher targeting in female are marked red. Green boxes highlight pathways associated with cell proliferation and purple boxes highlight pathways associated with immune response.

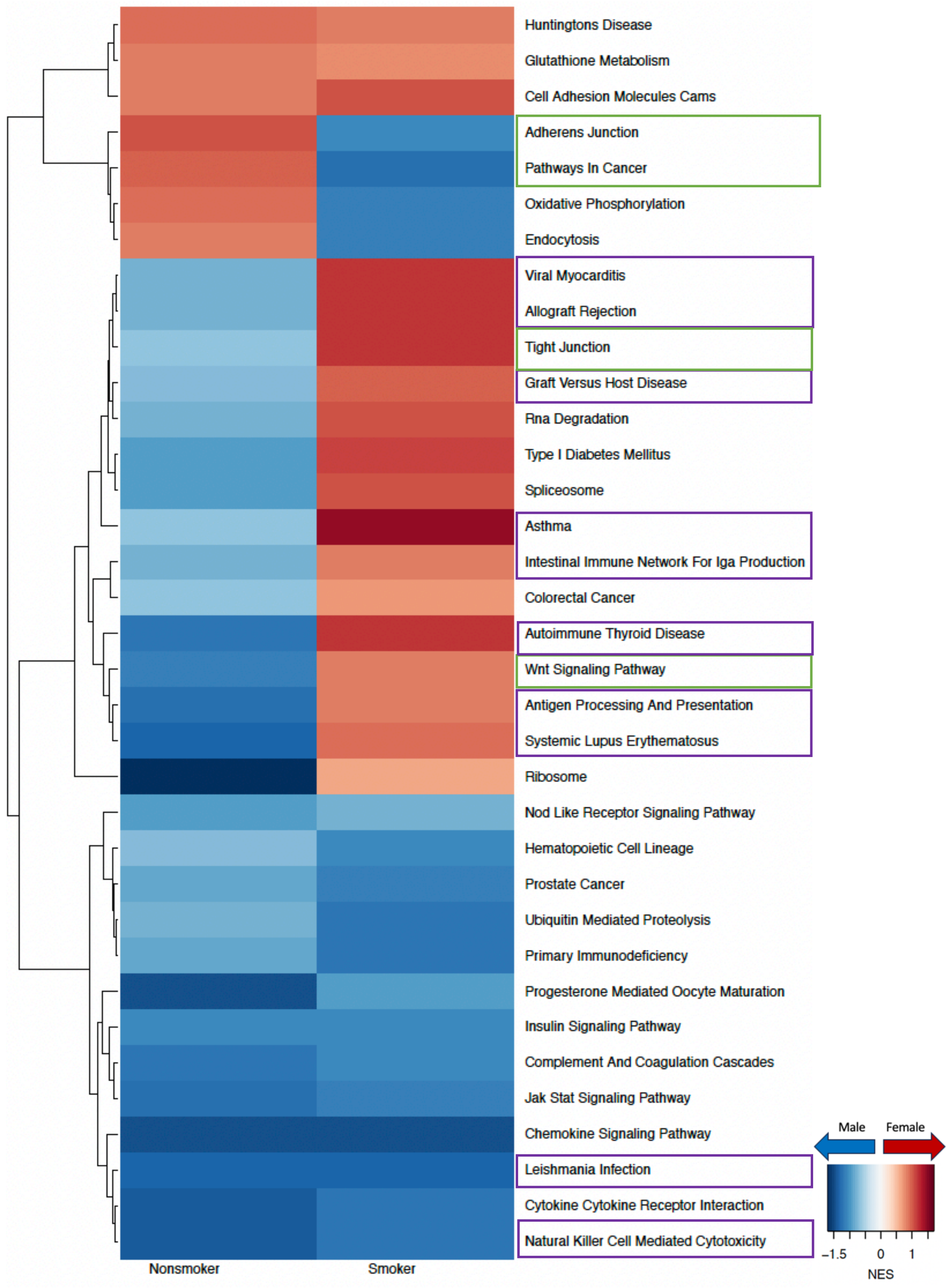

**Figure D.4**: Sex difference in immune and stromal cell composition in GTEx samples: nonsmokers (left) and smokers (right). Cell compositions are computed using “xcell”, which derives cell composition proportion of 36 immune and stromal, along with three composite scores: immune score, stroma score and microenvironment score. The bubbleplot shows only those cells that are significantly (p-value < 0.05) different in proportion in male and female samples.

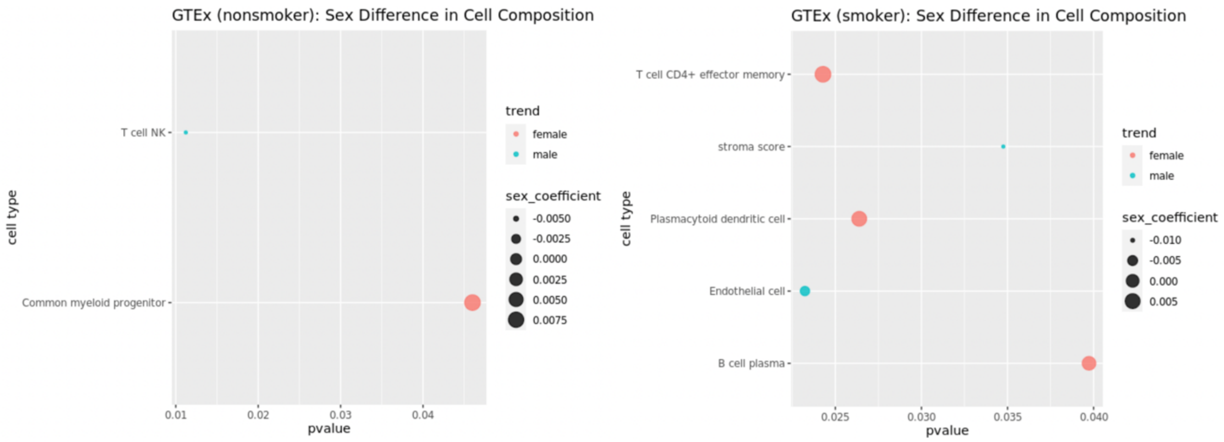


### E. Additional Tables

**Table E.1:** Distribution of Clinical Variables by Sex in GTEx.

| | MALE (N=395) | FEMALE (N=183) | P-value |
|---|---|---|---|
| **age** | | | |
| Mean (SD) | 54.1 (11.6) | 53.7 (12.4) | 0.767 |
| Median [Min, Max] | 57.0 [21.0, 70.0] | 55.0 [21.0, 70.0] | |
| **race** | | | |
| black or african american | 46 (11.6%) | 24 (13.1%) | 0.864 |
| others | 10 (2.5%) | 5 (2.7%) | |
| white | 339 (85.8%) | 154 (84.2%) | |
| **smoking_status** | | | |
| No | 107 (27.1%) | 73 (39.9%) | 0.00845 |
| Yes | 277 (70.1%) | 105 (57.4%) | |
| Unknown | 11 (2.8%) | 5 (2.7%) | |
| **ischemic_timeH** | | | |
| Mean (SD) | 8.31 (6.87) | 7.39 (7.18) | 0.146 |
| Median [Min, Max] | 7.12 [0, 24.4] | 6.10 [0, 24.4] | |

**Table E.2:** Distribution of Clinical Variables by Sex in TCGA.

| | MALE (N=238) | FEMALE (N=275) | P-value |
|---|---|---|---|
| **age** | | | |
| Mean (SD) | 65.4 (9.61) | 65.3 (10.1) | 0.853 |
| Median [Min, Max] | 66.0 [38.0, 88.0] | 66.0 [33.0, 87.0] | |
| **race** | | | |
| black or african american | 23 (9.7%) | 27 (9.8%) | 0.191 |
| others | 42 (17.6%) | 33 (12.0%) | |
| white | 173 (72.7%) | 215 (78.2%) | |
| **smoking_status** | | | |
| No | 20 (8.4%) | 55 (20.0%) | <0.001 |
| Yes | 211 (88.7%) | 213 (77.5%) | |
| Unknown | 7 (2.9%) | 7 (2.5%) | |
| **tumor_stage** | | | |
| stageI | 114 (47.9%) | 160 (58.2%) | 0.0917 |
| stageII | 67 (28.2%) | 54 (19.6%) | |
| stageIII | 38 (16.0%) | 46 (16.7%) | |
| stageIV | 14 (5.9%) | 12 (4.4%) | |
| not reported | 5 (2.1%) | 3 (1.1%) | |

**Table E.3:** Distribution of Clinical Variables by Sex in LGRC.

| | Male (N=49) | Female (N=59) | P-value |
|---|---|---|---|
| **age** | | | |
| Mean (SD) | 65.7 (10.8) | 61.9 (11.6) | 0.0817 |
| Median [Min, Max] | 67.0 [34.0, 87.0] | 63.0 [32.0, 82.0] | |
| **smoking** | | | |
| Never | 9 (18.4%) | 23 (39.0%) | 0.0332 |
| Ever | 36 (73.5%) | 29 (49.2%) | |
| NA | 4 (8.2%) | 7 (11.9%) | |

**Table E.4:** Distribution of Clinical Variables by Sex in GSE68465.

| | Male (N=218) | Female (N=214) | P-value |
|---|---|---|---|
| **age** | | | |
| Mean (SD) | 65.0 (9.42) | 64.0 (10.7) | 0.272 |
| Median [Min, Max] | 65.5 [35.0, 82.0] | 64.0 [33.0, 87.0] | |
| **tumor_stage** | | | |
| 1 | 65 (29.8%) | 83 (38.8%) | 0.0717 |
| 2 | 126 (57.8%) | 116 (54.2%) | |
| 3 | 20 (9.2%) | 8 (3.7%) | |
| 4 | 6 (2.8%) | 6 (2.8%) | |
| NA | 1 (0.5%) | 1 (0.5%) | |
| **race** | | | |
| White | 125 (57.3%) | 164 (76.6%) | <0.001 |
| Black or African American | 6 (2.8%) | 6 (2.8%) | |
| others | 4 (1.8%) | 2 (0.9%) | |
| Unknown | 83 (38.1%) | 42 (19.6%) | |
| **smoking** | | | |
| Never | 9 (4.1%) | 39 (18.2%) | <0.001 |
| Ever | 148 (67.9%) | 142 (66.4%) | |
| Unknown | 61 (28.0%) | 33 (15.4%) | |
